# Nucleolar structure connects with global nuclear organization

**DOI:** 10.1101/2023.03.30.534966

**Authors:** Chen Wang, Hanhui Ma, Susan J. Baserga, Thoru Pederson, Sui Huang

## Abstract

The nucleolus is a multi-functional nuclear body. To tease out the roles of nucleolar structure without resorting to multi-action drugs, we knocked down RNA polymerase I subunit RPA194 in HeLa cells by siRNA. Loss of RPA194 resulted in nucleolar structural segregation and effects on both nucleolus-proximal and distal nuclear components. The perinucleolar compartment was disrupted, centromere-nucleolus interactions were significantly reduced, and the intranuclear locations of specific genomic loci were altered. Moreover, Cajal bodies, distal from nucleoli, underwent morphological and compositional changes. To distinguish whether these global reorganizations are the results of nucleolar structural disruption or inhibition of ribosome synthesis, the pre-ribosomal RNA processing factor, UTP4, was also knocked down, which did not lead to nucleolar segregation, nor the intranuclear effects seen with RPA195A knockdown, demonstrating that they do not arise from a cessation of ribosome synthesis. These findings point to a commutative system that links nucleolar structure to the maintenance and spatial organization of certain nuclear bodies and genomic loci.

## Introduction

The nucleus is highly compartmentalized, with interphase chromosomes organized into specific territories and interchromatin space occupied by various non-membrane enclosed organelles, such as Cajal and PML bodies, and interchromatin granule clusters (Misteli and Spector, 2011; Pederson, 2011). The nucleolus is the most prominent nuclear body and is the site of ribosome synthesis as well as other functions that have since come into view, including assembly of the signal recognition particle, cell cycle progression, macromolecular trafficking, and surveillance of genotoxic and nucleolar stress (Baserga and DiMario, 2018; Baserga et al., 2020; Lin et al., 2014; Nemeth and Grummt, 2018; Pederson, 2011; Pederson and Tsai, 2009). Here we have deployed a new strategy to explore whether the nucleolar structure might have a additional role in the cell that are independent from ribosome synthesis.

A frequently deployed tool for investigating the nucleolus has been exposure of cells to a low concentration of actinomycin D. A pioneering study that correlated a reduction of ^3^H-uridine labeling of the nucleolus with a biochemical demonstration of impaired ribosomes synthesis (Perry, 1962) was one of the key steps in defining the nucleolus as the site of ribosome biogenesis. It was subsequently found that exposure of mammalian cells to low concentrations of actinomycin D also leads to alterations of nucleolar morphology, in which its three ultrastructurally-defined components, fibrillar center (FC), dense fibrillar components (DFC) and granular components (GC), undergo repositioning, (Clark et al., 1967). At these low concentrations, actinomycin D preferentially intercalates and alkylates DNA at adjacent GC base pairs (Gallego et al., 1997; Sengupta et al., 1988). Since ribosomal RNA genes of eukaryotic cells are GC-rich (at least 65-70% in the case of most mammalian genomes), there has been a tendency to regard low actinomycin as a highly specific experimental intervention. However, there are numerous other GC-rich sites throughout mammalian genomes. Indeed, our bioinformatics survey of the human reference genome revealed 320 sites which are 65-70% GC, including some that are 85% or more (Table S1). Thus, it is possible that nucleolar disruption by actinomycin D and concurrently observed effects could be attributable to the drug’s action at one or more of these hundreds of genomic sites or to other yet to be defined actions of the drug.

The circumvent the pitfalls of using small molecule drugs with multi-valent functions, we employed siRNA approach to disrupt rDNA transcription machinery to determine whether the loss of transcription machinery impacts the well-defined nucleolar structure as actinomycin D treatment does. We also address whether nucleolar structure maintenance is dependent upon ribosome synthesis and whether nucleolar structural integrity plays a role in the organization of other nuclear domains.

## Results

### Knockdown of RPA194 phenocopies low actinomycin D treatment

We used siRNA knockdown of RPA194, the largest subunit of RNA polymerase I (Russell and Zomerdijk, 2005), to disrupt rDNA transcription. Seventy-two hours after transfection, the level of RPA194 was reduced by more than 95% while the levels of another rDNA transcription factor, UBF, remain unchanged (Fig. 1A). Knockdown of RPA194 led to an inhibition of rRNA synthesis as measured both by pulse-labeling cells with BrU and qRT-PCR of the 5ʹ ETS regions of the nascent pre-rRNA (Suppl. Fig. 1A and 1B). As expected, there was a resultant reduction of ribosome synthesis as assessed by tracking an inducible GFP-tagged ribosomal protein RPL29 (Wild et al., 2010) (Fig. 1B). In untreated and control siRNA transfected cells, GFP-RPL29 was localized in both nucleoli and the cytoplasm within 24 hours of activation, reflecting its assembly into new ribosomes and localization to the cytoplasm. In comparison, in RPA194 knockdown cells, GFP-RPL29 was detected only in nucleoli and not in the cytoplasm 24 hours after activation, indicating the lack of newly synthesized ribosomal particles bearing GFP tagged RPL29 entering the cytoplasm. When the pre-ribosomal RNA processing factor UTP4 (Freed et al., 2012) was knocked down, ribosome synthesis was similarly blocked (Fig. 1B), as also represented by a similar lack of cytoplasmic GFP-RPL29 signal. These results demonstrate the inhibition of ribosome synthesis by knockdown of either RPA194 or UTP4.

**Figure 1:**
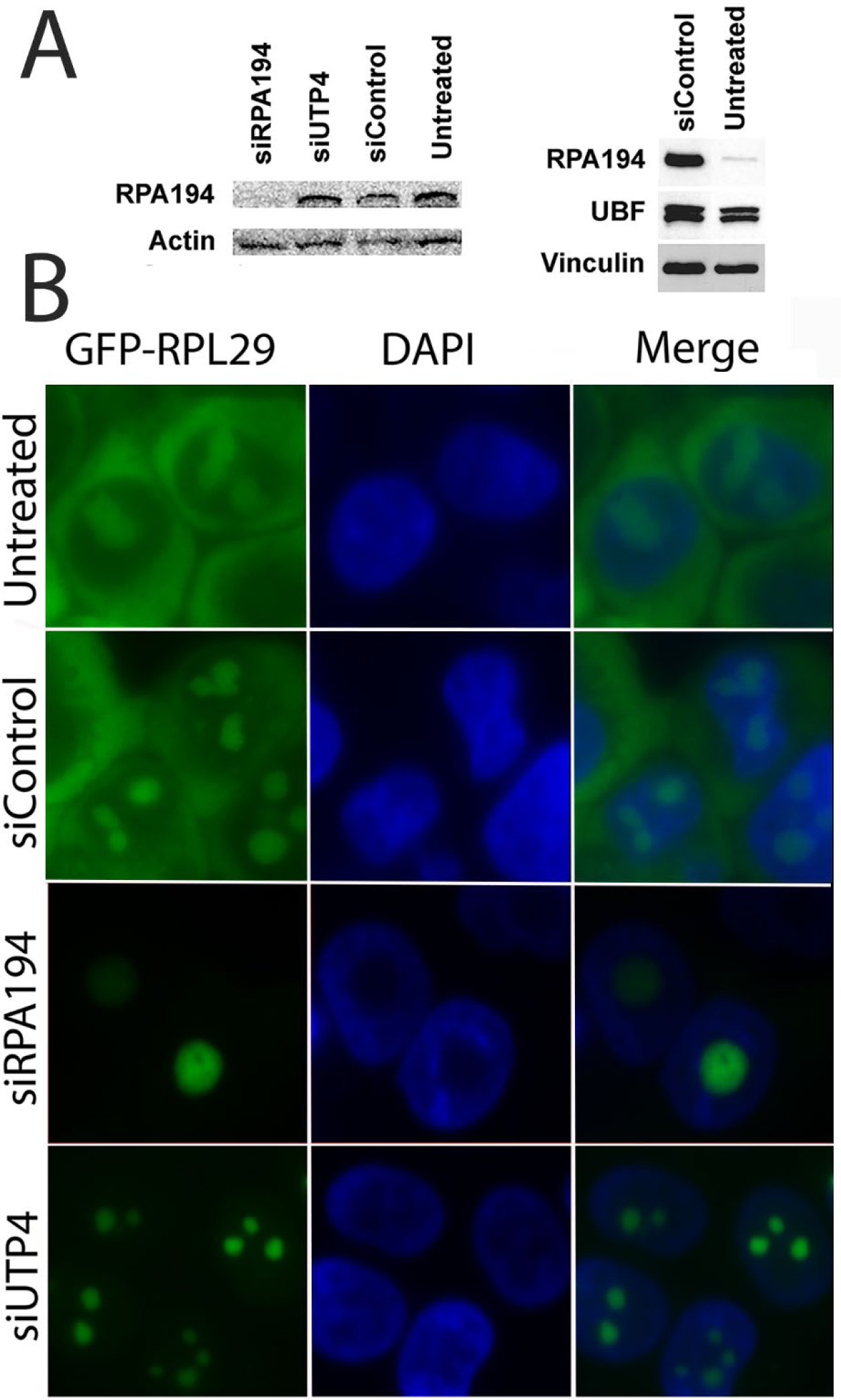
siRPA194 and siUTP treatment in HeLa cells significantly reduced ribosome synthesis, but only siRPA194 treatment induces nucleolar segregation. (A) The level of RPA194 protein was significantly reduced while the UBF level was unchanged at 72 hours after transfection. (B) Both siRPA194 and siUTP4 treatment inhibited ribosome synthesis as assayed in an inducible GFP-RPL29 stably transfected HeLa cell line. Activation of fusion protein expression 48 hours after transfection showed little presence of GFP-RPL29 in the cytoplasm in siRPA194 and siUTP4 compared to control cells, demonstrating lack of new ribosome synthesis. Bars = 5 mm

Electron microscopy (EM) revealed that knockdown of RPA194 resulted in a reorganization of nucleolar architecture. The nucleolus normally contains three distinct ultrastructural components: FC (Fig. 2A, arrows), DFC (Fig. 2A, arrowheads) and GC (Fig. 2A, asterisk) (Pederson, 2011; Raska, 2003). Knockdown of RPA194 induced segregation of the fibrillar components from the granular components (Fig. 2A, left panel). The nucleolar segregation observed by EM corresponds to the redistribution of the fibrillar protein, UBF (Fig. 2B, arrowheads) and the dense fibrillar component marker protein fibrillarin (Suppl. Fig. 2) from their normal nucleolar location into nucleolar caps (Fig. 2B, arrows). Quantitative analyses revealed that the loss of RPA194 in siRNA treated cells closely paralleled nucleolar segregation (Fig. 2C), similar to that observed in cells treated with low concentrations of actinomycin D (Suppl. Fig. 3). In comparison to the knockdown of RPA194, siUTP4 treatment, while also blocking ribosome synthesis, did not lead to nucleolar segregation (Fig. 1B, 2A,B and C). This demonstrated, for the first time, that nucleolar segregation is not the result of ribosome synthesis inhibition, but rather the disruption of rDNA transcription machinery.

**Figure 2:**
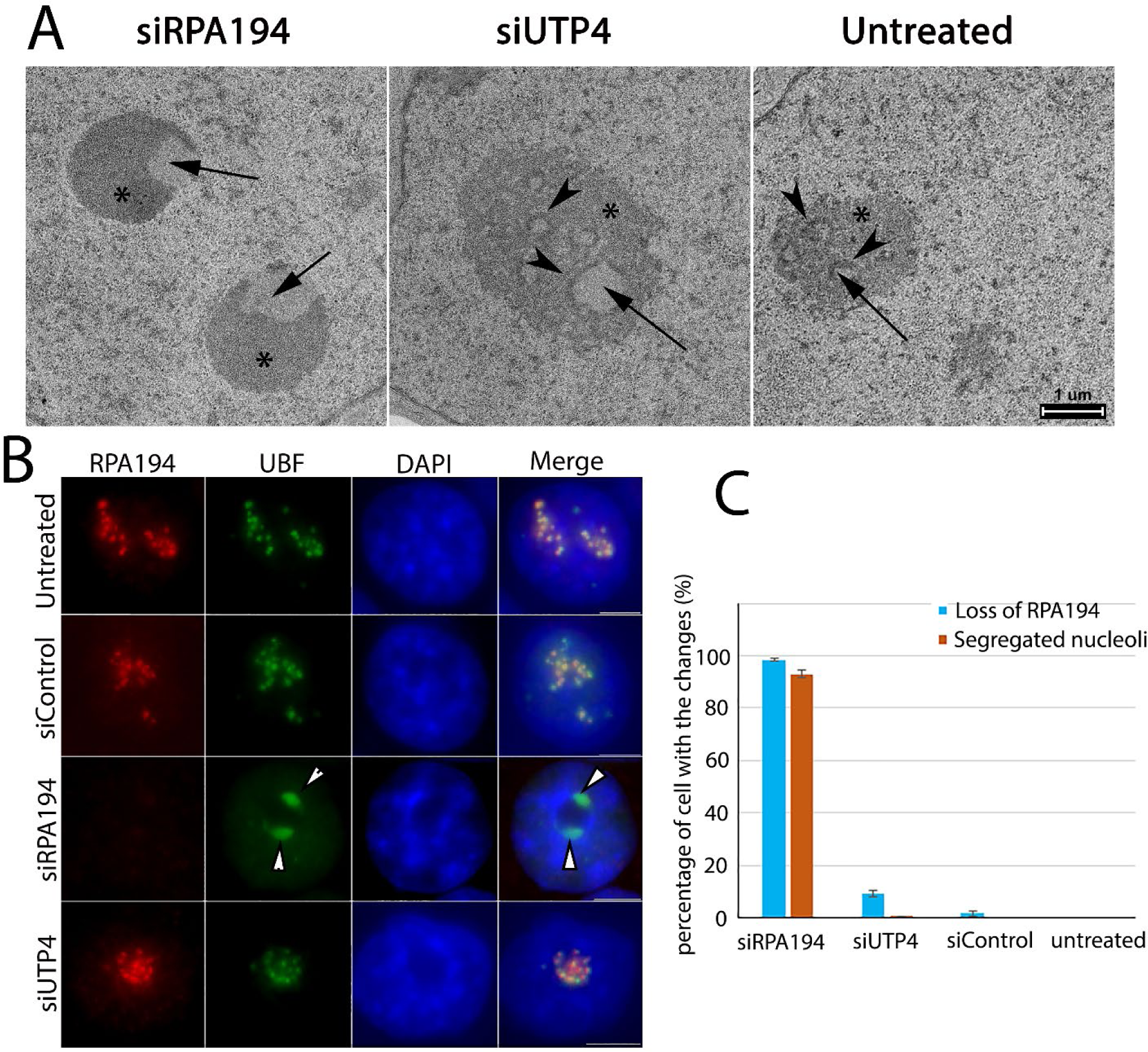
siRPA194 knockdown cells induced nucleolar segregation. (A) Examination by electron microscopy demonstrated the siRPA194 treatment (left panel) in HeLa cells changed nucleolar structures where the fibrillar components (arrows) were segregated from the granular components (*). In comparison, the nucleolus in UTP4 knockdown cells (middle panel) maintained the typical three structural components, fibrillar center (arrow), dense fibrillar components (arrowhead), and granular components (*) as those in the siControl or untreated cells (right panels). (B) Correspondingly, siRPA194 treated, but not siUTP4 treated or control cells, showed segregated caps like localization of UBF as detected at the immunofluorescence microscope level. (C) Quantification showed that the segregation of nucleoli occurs mostly in RPA194 knockdown HeLa cells (n>100 cells, p<0.005 between siRPA194 and all other conditions). Bars =5 mm.

The spatial redistribution of the fibrillar centers upon nucleolar segregation implies that the rDNA does so in parallel. To determine if this was the case, we deployed CRISPR-dCas9 mediated labeling of rDNA clusters in siRNA treated and control cells (Ma et al., 2015; Ma et al., 2019; Ma et al., 2018; Ma et al., 2016a; Ma et al., 2016b). Sixteen sets of CRISPR-Cas9 gs RNA were designed to label rDNA. The specificity of the labeling was validated by simultaneous labeling with an anti-UBF antibody on mitotic Nucleolar Organizer Regions (NORs) (Suppl. Fig. 4) as it has been shown that rDNA transcription factors remain bound to active NORs throughout mitosis (Roussel et al., 1996). In untreated cells, control siRNA or siUTP4 transfected cells, the labeled rDNA was localized in the nucleoli in the expected nucleolar pattern. But after RPA194 knockdown, the rDNA signals became redistributed and colocalized with the repositioned fibrillarin caps (Fig. 3A and C). This was also the case with low actinomycin treatment (Fig. 3B and C), consistent with findings from a recent study (Mangan and McStay, 2021). In both cases, nearly all the rDNA was associated with the fibrillar protein-localized caps (Fig. 3A, B, and C). These findings together with those in Fig. 1 show that RPA194 knockdown recapitulates in the classical phenomenon of nucleolar segregation observed with the treatment of low actinomycin.

**Figure 3:**
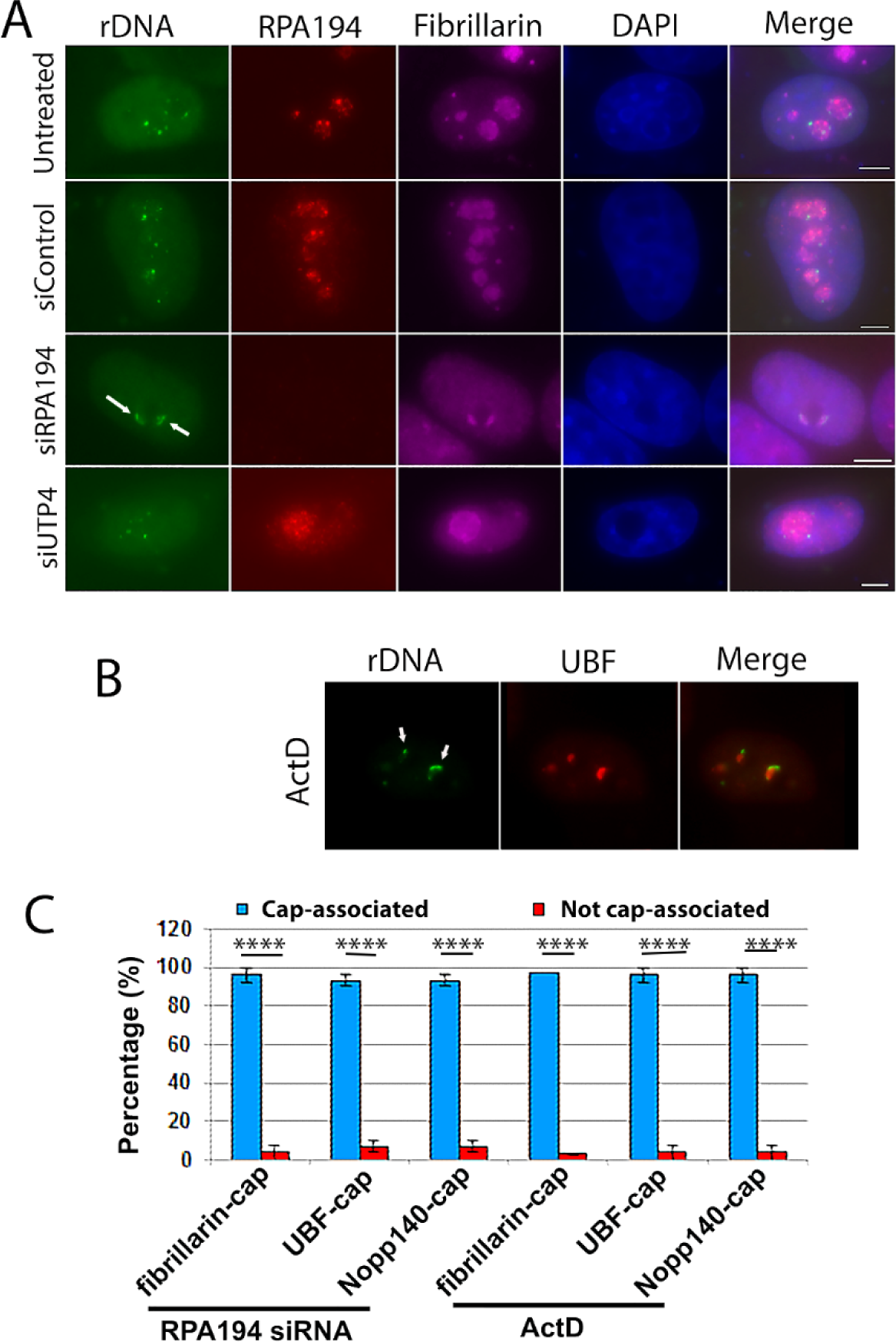
rDNA co-segregated with the fibrillar proteins (fibrillarin or UBF) present in cap structures in segregated nucleoli. (A) Eleven sets of CRISPR-Cas9 sgRNA specifically complementing the rDNA were labeled with MS2-GFP, showing the localization of the rDNA chromatin, which redistributed with the fibrillar components to the nucleolar caps in siRPA194 treated HeLa cells (arrows) similarly to (B) those treated by ActD (arrows). (C) Quantitative evaluation indicated the co-partitioning of rDNA chromatin with the fibrillar components in nearly all of cells treated by siRPA194 or ActD (n>100 cells). Bars = 5 mm

### Nucleolus-proximal nuclear domains are altered by knockdown of RPA194, but not UTP4

With siRPA194 knockdown as a selective tool to disrupt nucleolar structure without the drawbacks of drug treatment, we asked whether nucleolar structure might play a role in nuclear organization. The perinucleolar compartment (PNC) is a distinct structure in many cancer cell types, including HeLa cells, and is physically associated with the nucleolus (Norton et al., 2008; Pollock and Huang, 2009). In a previous study, knockdown of RPA194 resulted in disassembly of PNCs (Frankowski et al., 2018). To investigate whether PNC disruption is the consequence of ribosome synthesis or rather due to nucleolar structure alteration, we included siUTP4 knockdown as a control. Antibodies against PTB (polypyrimidine binding protein, a fiduciary marker protein of the PNC) were used to immunolabel PNCs. As shown in Fig. 4A and B, while the siRPA194 treatment reduced PNC prevalence, knockdown of UTP4 did not significantly impact PNC bodies. Additionally, most of the remaining PNCs underwent structural alteration, becoming crescent moon shaped (Fig. 4A, arrows) in siRPA194 treated cells. Thus, PNC integrity is dependent on the nucleolar structure or rDNA transcription machinery, rather than ribosome synthesis.

**Figure 4:**
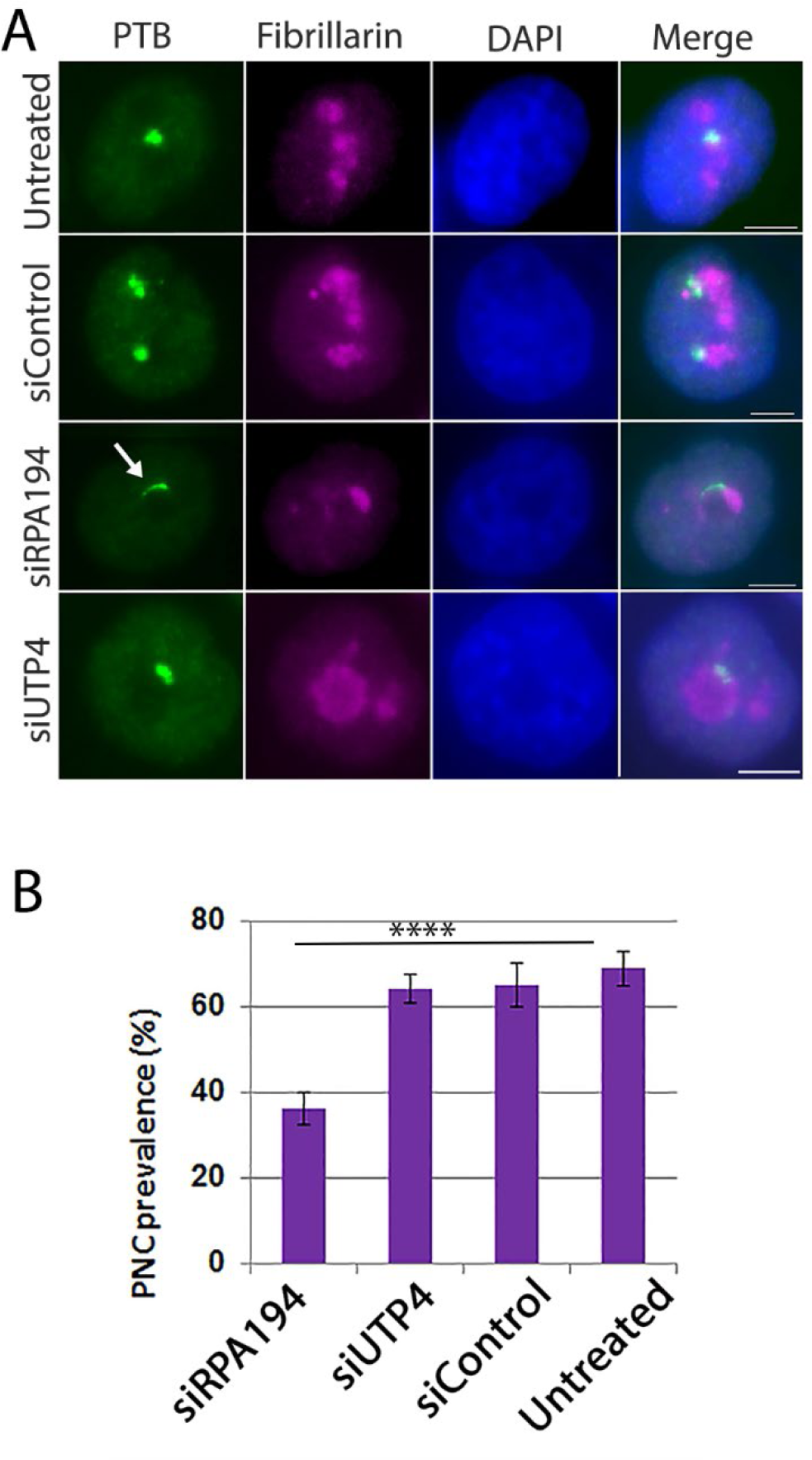
PNC structure and prevalence were altered in siRPA194, but not in siUTP4 treated HeLa cells. (A) The shape of most PNCs was changed, becoming extended (arrows) in siRPA194 treated, but not in siUTP4 treated or control cells. (B) Quantitative analyses demonstrated a significant reduction of PNC prevalence in siRPA194 treated cells (n>100 cells). Bars = 5 mm ****p<0.001

Centromeres are known to cluster around nucleoli (Padeken and Heun, 2013), with an average 60% of the centromeres situated close to nucleoli in the HeLa cells we have used here (Foltz et al., 2009). To evaluate whether nucleolar segregation impacts nucleolus-centromere proximity, we immunolabeled centromeres (of all chromosomes) and nucleoli with anti-CENPA and anti-UBF antibodies respectively. As shown in Figure 5A and B, upon RPA194 knockdown centromeres were less frequently associated with nucleoli and more dispersed throughout the nucleus. In contrast, UTP4 knockdown did not change nucleolus-centromere associations (Fig. 5A and B). These findings indicate that the nucleolus-proximal centromere arrangement is contingent on normal nucleolar organization or Pol I transcription.

**Figure 5:**
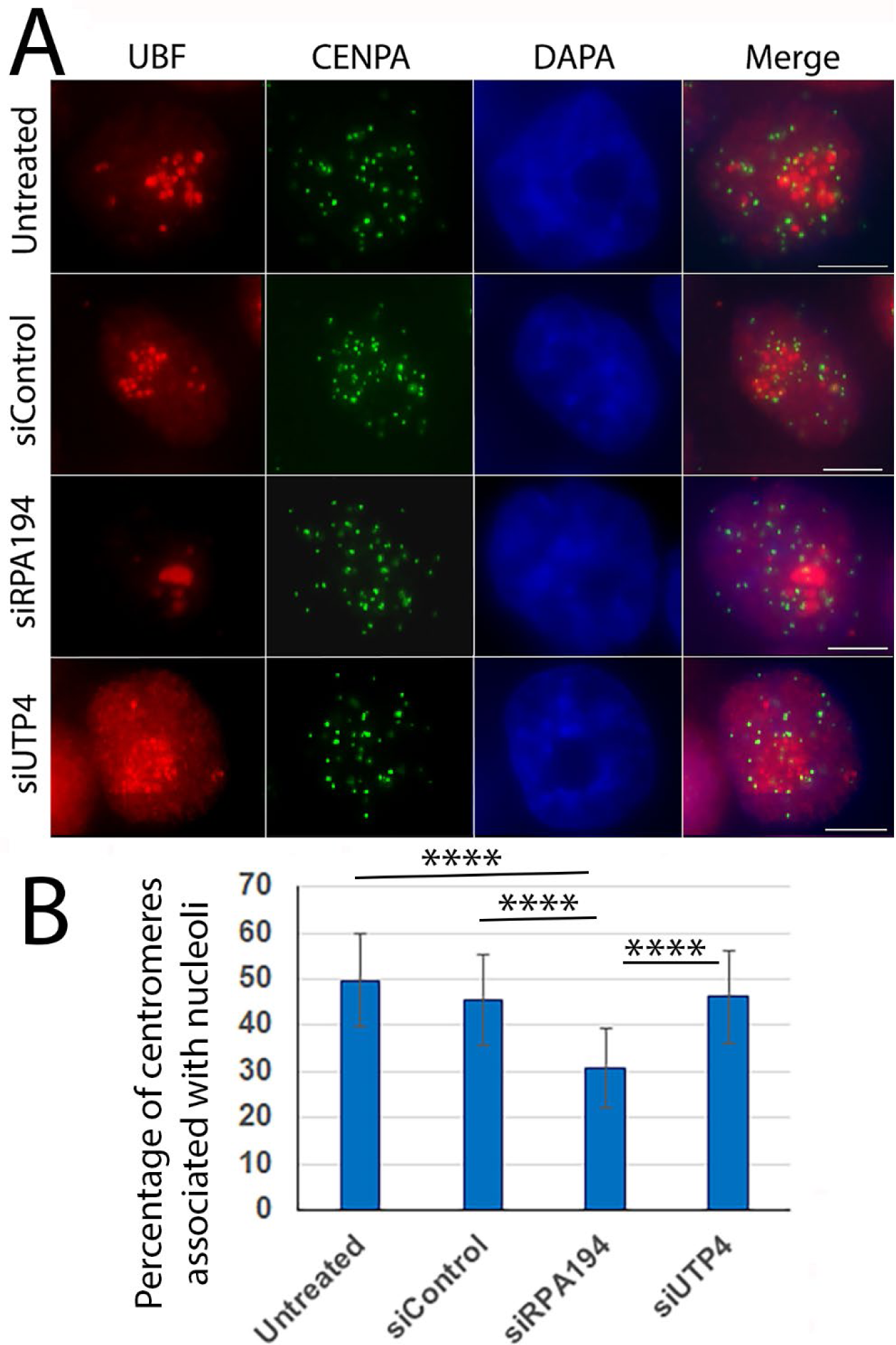
RPA194 knockdown changed nucleoli-centromere interactions (A) The clustering of centromeres (green) around nucleoli (red) decreased in siRPA194 treated, but not in siUTP4 treated and control cells. (B) Quantifying analyses showed changes were statistically significant (n>50 cells) ****p<0,001. Bars = 5 mm

### Repositioning of genomic loci after RPA194 knockdown

We next asked whether there are more wide-ranging effects throughout the 3-D nucleome upon RPA194 knockdown and the resulting nucleolar reorganization. As shown in Fig. 6A, the HIST1H2BN locus underwent a repositioning from its mainly nucleolar proximity to a predominantly central nuclear location in siRPA194 treated cells. Conversely, the predominantly central MALL locus become repositioned to more nuclear-peripheral positions (Fig. 6B). These positional changes of genomic loci were not observed in either siUTP4 or control oligo treated cells, nor in untreated cells, and thus are specifically attributable to RPA194 knockdown coupled nucleolar segregation.

**Figure 6:**
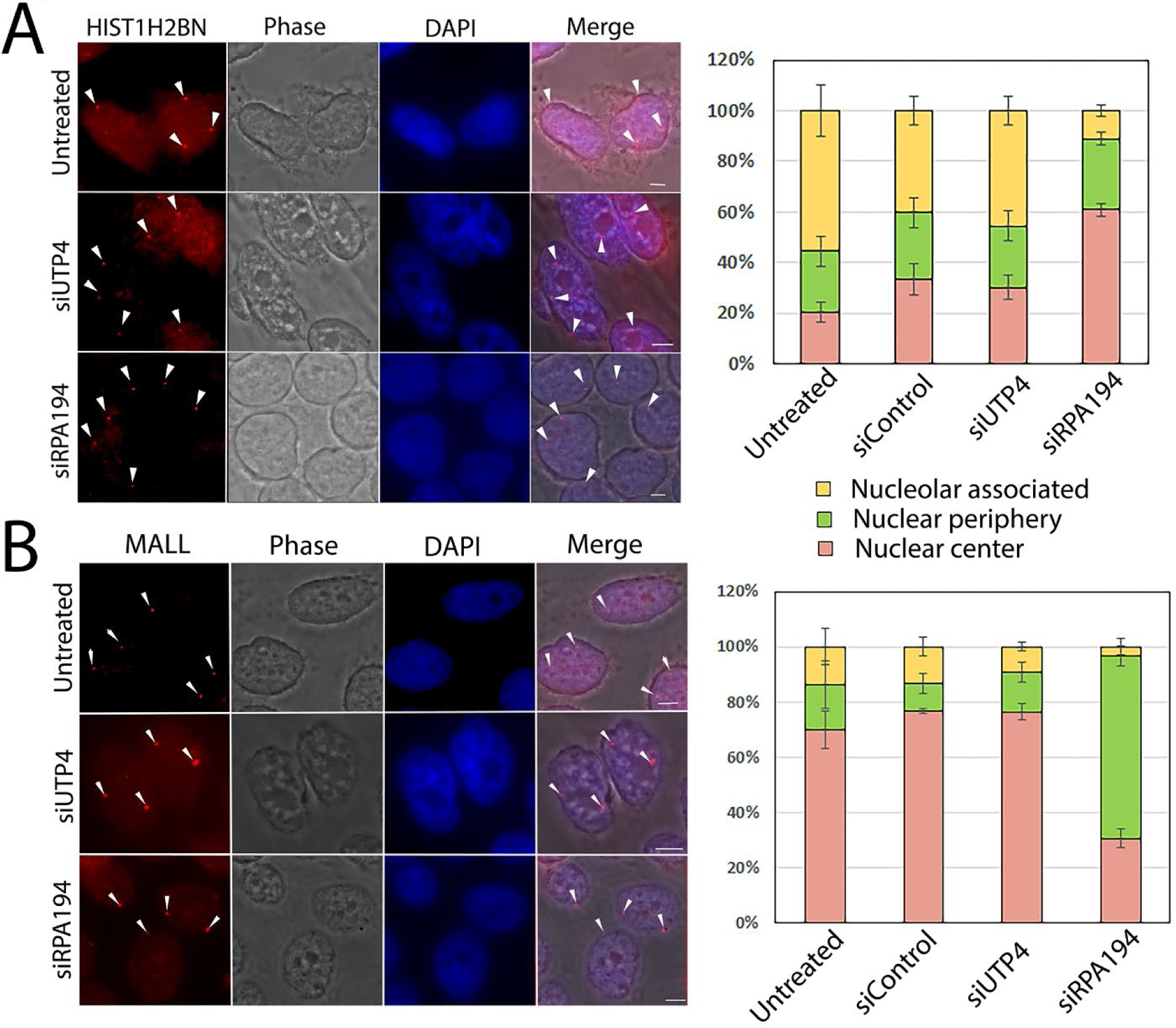
RPA194 knockdown changed spatial distributions of some genes. (A) HIST1H2BN gene loci became more centrally localized in siRPA194 treated HeLa cell nuclei than siUTP4 treated and other control cells (arrowheads) whereas (B) MALL loci became more nuclear peripherally localized in siRPA194 treated cells (arrowheads). Corresponding quantitative analyses showed significant differences in central and peripheral localization of the gene loci between treated and control cells. (n>50 cells) p<0.01. Bars = 5 mm

### Nucleolar organization governs Cajal body integrity

Given these results, we wondered if normal nucleolar organization might play a role not only in the intranuclear spatial location of certain genomic loci, but perhaps also in the maintenance of nuclear bodies. We therefore examined Cajal bodies (Gall, 2003; Nizami et al., 2010; Verheggen et al., 2002; Wang et al., 2016) and found that RPA194 knockdown, but not siUTP4 knockdown, led to their disruption as tracked both by the localization of p80 coilin, a Cajal body marker, and fibrillarin or UBF (Fig. 7A, and Suppl. Fig. 5). The disruption of Cajal bodies was closely correlated with nucleolar segregation (Fig. 7B) like those previously described in actinomycin D-treated cells (Shav-Tal et al., 2005). The dispersion of Cajal bodies was also observed when examined with the additional marker protein SMN (Fig. 8). In addition to the dramatic structural changes, the Cajal body proteins, SMN and coilin no longer perfectly colocalized as shown in Fig. 8. These findings demonstrate that the nucleolus, in its normal operation, not only impacts genomic organization, but also a class of nuclear bodies usually situated distant from it.

**Figure 7:**
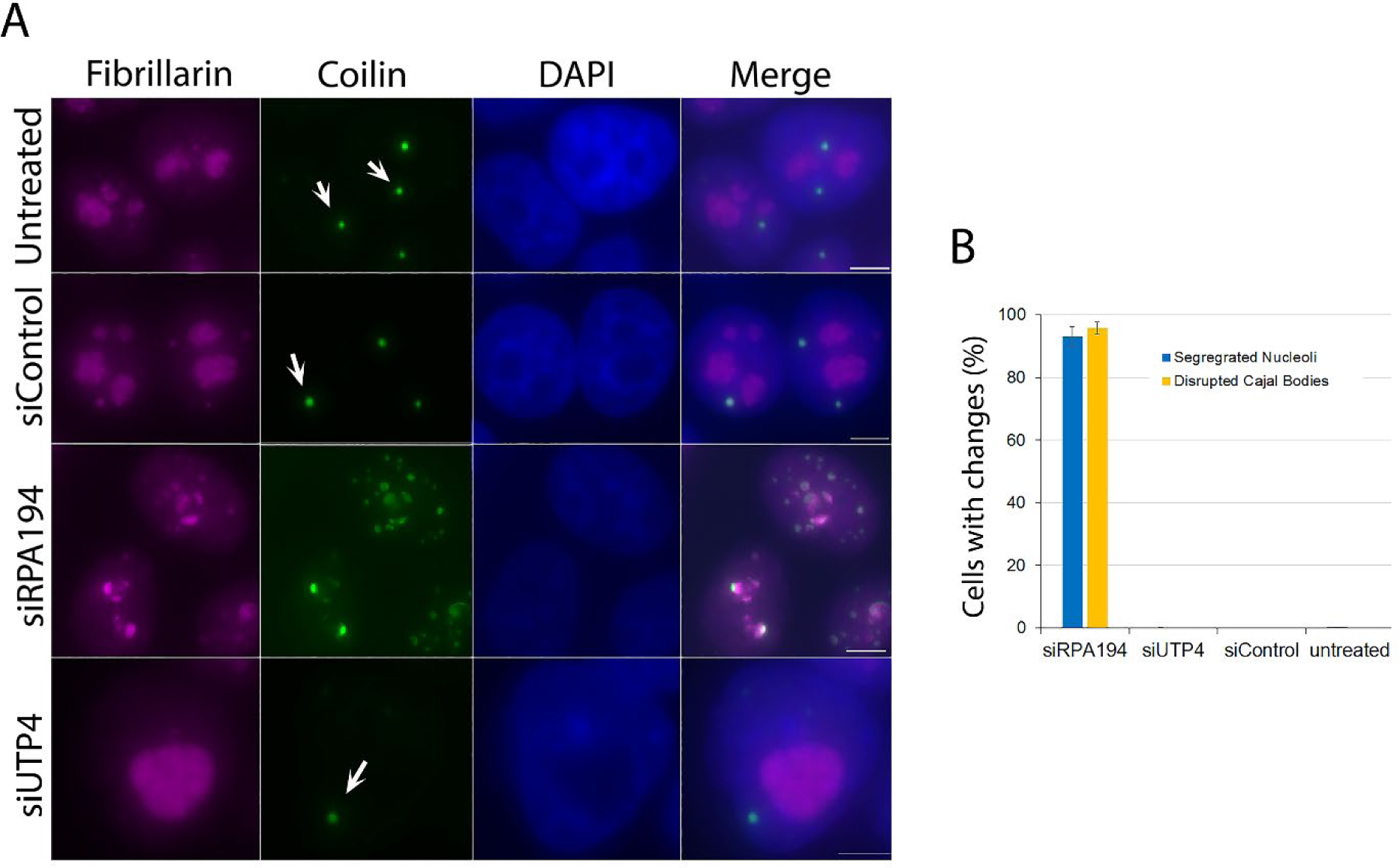
siRPA194, but not siUTP4 treatment, induced disruption of Cajal bodies (arrows). (A) Upon siRPA194 treatment, Cajal bodies became irregularly shaped and some juxtaposed segregated nucleoli in siRPA194 treated cells (third panels), as detected using anti-coilin antibodies. However, Cajal bodies remained round-punctuated spots (arrows) in siUTP4 or control treated cells. (B) Quantitative evaluation showed a close coupling between nucleolar segregation and disruption of Cajal bodies (n>100 cells) (p<0.001 between siRPA194 and all others).

**Figure 8:**
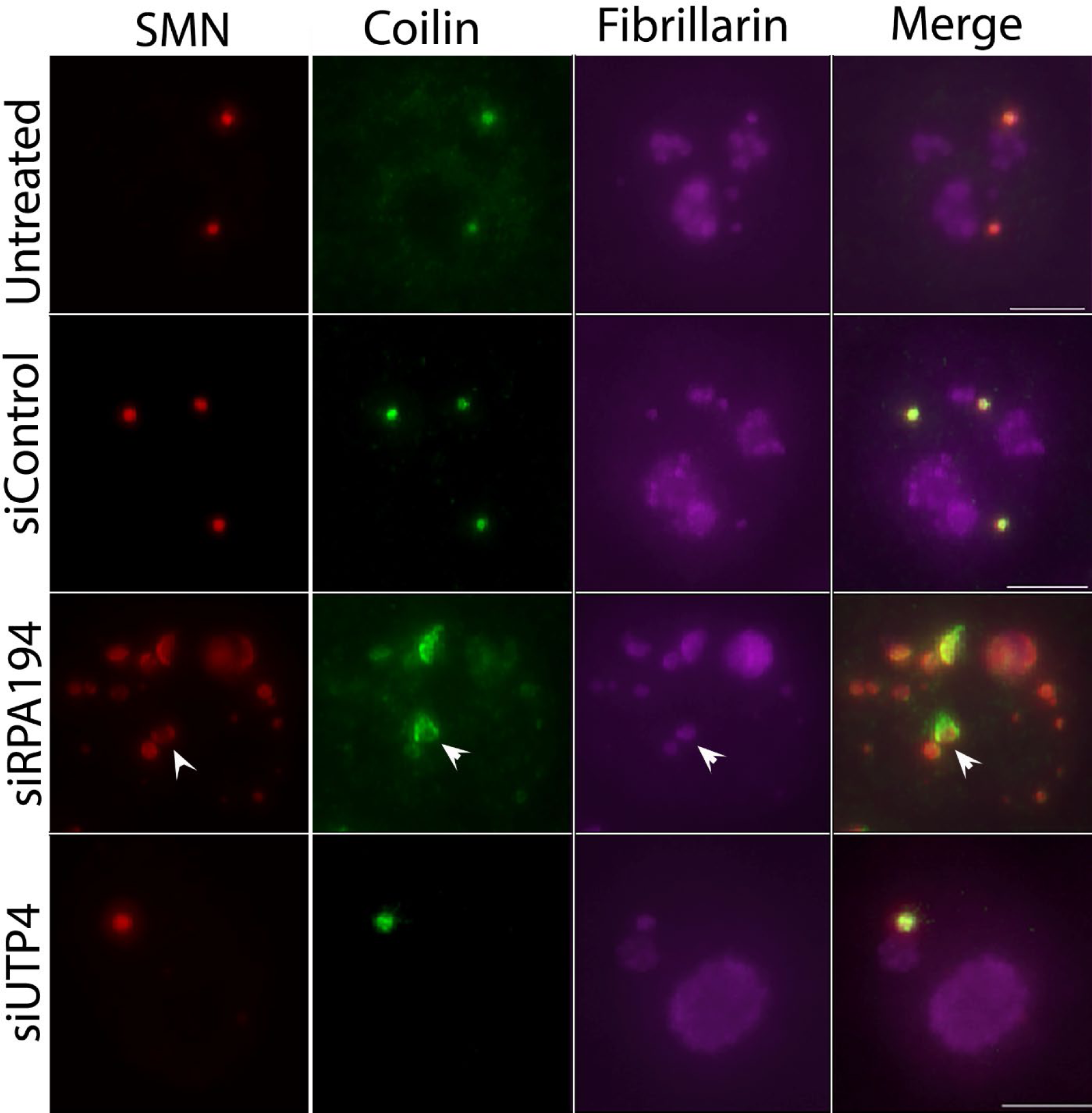
Cajal body components no longer showed nearly complete colocalization in cells with segregated nucleoli from the treatment of siRPA194. Each panel shows a single nucleus. SMN and coilin which normally colocalize to the dotted-like Cajal body structures (Merge) no longer perfectly coincide in the round spots, rather become less colocalized in siRPA194 treated cells (arrowhead). Bar= 5mm

The observed disruption of Cajal bodies prompted us to ask if RPA194 knockdown influences another nuclear body, the histone locus body (Nizami et al., 2010). As shown in Suppl. Fig. 6, there was no effect of RPA194 knockdown on histone locus bodies, demonstrating that not all nuclear bodies are regulated by an intact nucleolar structure.

## Discussion

To circumvent the lack of specificity of low actinomycin D treatment as a tool to investigate the possible roles of nucleolar structure in global nuclear organization, we deployed knockdown of the Pol I subunit, RPA194, and of the pre-rRNA processing factor, UTP4, both leading to ribosome synthesis inhibition. However, only siRPA194 knockdown, but not UTP4 knockdown, induced nucleolar segregation, indicating that continuous ribosome synthesis is not required for maintenance of the classical tripartite nucleolar ultrastructure and gave us an opportunity to specifically address the impact of nucleolar structure on other nuclear components. We chose to use HeLa cells because they lack the p53 protein, which might otherwise trigger apoptosis upon RPA194 knockdown. We found that nucleolar segregation leads to changes throughout the 3D nucleome, in which nucleolus-proximal entities such as the PNC and centromere clusters reorganize significantly, certain genomic loci change their spatial distribution from nucleolus-proximal to central nuclear locations or from mostly nuclear central to the nuclear periphery. In addition, nucleolar segregation disrupts nucleoplasmic Cajal bodies, but not histone locus bodies.

Our findings demonstrate a connection between nucleolar structure and a subset of various intranuclear components or genomic loci, with no apparent connection with others. The nature of the underlying signaling network is not clear. One possibility is that nucleolar segregation alters physical contacts between the nucleolar surface and directly connecting elements, such as nucleolus-proximal heterochromatin (Vertii et al., 2019), other nucleolus-associated domains of the genome (Nemeth et al., 2010; van Koningsbruggen et al., 2010), PNCs, or clustered centromeres. There is, moreover, extensive evidence for shuttling of proteins between the nucleolus and the nucleoplasm both in normal circumstances (Chen and Huang, 2001; Phair and Misteli, 2000) and in actinomycin D-induced nucleolar segregation (Shav-Tal et al., 2005). Thus, it is also possible that nucleolar segregation restricts the dynamics and exchanges of nucleolar and nuclear components that are part of the regulatory network important to nuclear homeostasis. Future studies will investigate the mechanics and mechanisms of these connections.

## Material and Methods

### Cell culture

HeLa cells were maintained in DMEM (<u>ThermoFisher Scientific</u>) with 10% FBS (GEMINI Bio Products) and 100 units/ml of penicillin and streptomycin (<u>ThermoFisher</u> <u>Scientific</u>). RPL29-GFP cells were cultured in DMEM with 0.3mg/ml of hygromycin and 0.5ug/ml puromycin. Cells were generally plated 24 h prior to treatment with siRNA.

### RNA interference

The siRNA targeting human RPA194 was obtained from Invitrogen (cat#: 10620318) as was a negative control siRNA (cat#: 12935-300). The siRNA targeting human UTP4 was obtained from GE Dharmacon (cat#: M-015011-01-0005). siRNA were transfected into cells with Lipofectamine® RNAiMAX Transfection Reagent (Thermofisher Scientific; cat#: 13778) according to manufacturer’s instructions. The reported experiments were performed after 72 hours transfection.

### Immunofluorescence

Primary antibodies were from the indicated sources and used at the stated dilutions: RPA194, Santa Cruz Biotechnology, cat #sc-48385, 1:100; PTB (S. Huang laboratory), 1:300; UBF, Santa Cruz Biotechnology, cat#: sc-13125, 1:50; fibrillarin, Sigma, cat #: ANA-N, 1:10; CENPA (Thermo Fisher #MA1-20832, 1:100; coilin, Santa Cruz Biotechnology, cat#: sc-32860, 1:300; Flash histone locus antibody protein (Gall lab), 1:500; SMN, Santa Cruz Biotechnology, cat.# sc-15320, 1:300. The fluorescein conjugated (Jackson ImmunoResearch) and Alexa Fluor-labeled (ThermoFisher) secondary antibodies were used at a dilution of 1:200. Slides were visualized on a Nikon Eclipse Ti-E inverted fluorescence microscope using NIS-Elements AR 4.2 software (Nikon).

### Western blots

These were performed according to manufacture protocols (Millipore). The primary antibodies and dilutions used were RPA194 (see above), 1:500); UBF (see above), 1:500); beta-actin (Sigma, cat.# A5060), 1:3000, as loading controls. LI-COR IRDye secondary antibodies (1:10,000) were used for visualization. Protein bands were detected using LI-COR Odyssey Image Studio (LI-COR Biosciences).

### Electron microscopy

Cells were in fixed 2% paraformaldehyde-2.5% glutaraldehyde in 0.1M cacodylate buffer, pH 7.3, processed and examined in the Northwestern University Feinberg School of Medicine Imaging Facility.

### Inducible expression of RPL29-GFP HeLa cell line

The tetracycline-inducible RPL29-GFP cell line was maintained in hygromycin and puromycin. Before each experiment the cells were transferred to medium without the antibiotics and transfected with the desired siRNA or left untreated. After 48 hours, tetracycline was added at 125 ng/ml to induce RPL29-GFP expression followed by examination 20 hours later.

### In situ hybridization

HeLa cells were seeded to 6 well plates one day before transfection of siRNAs. Three days post-transfection, the genomic loci HIST1H2BN and MALL were detected using BACs obtained from bac.chori.org (cat. #’s RP-11-958P15 and RP11-349F11, respectively, and labeled by nick translation Abbott Laboratories (cat. # 07J00-001. The hybridization signals were measured using the Z series photos taken on a Nikon Eclipse Ti-E inverted fluorescence microscope. The gene loci spatial positions were analyzed in their relation to the nuclear envelope and nucleoli, and the space between.

### CRISPR dCas9 labeling of rDNA

The CRISPR Cas9 rDNA labeling constructs were “Nm dCas9 3XGFP” as detailed in (Ma et al., 2015) and a U6 promoter guide RNA plasmid containing a pool of 16 fragments complementary to rDNA. The sequences are detailed here. rDNA-F1: ACCGCGCGTGTTCTCATCTAGAAG; rDNA-R1: AAACCTTCTAGATGAGAACACGCG; rDNA-F2: ACCGTGGGCAGAACGAGGGGGACC; rDNA-R2: AAACGGTCCCCCTCGTTCTGCCCA; rDNA-F3: ACCGCGGACACAGCACTGACTACC; rDNA-R3: AAACGGTAGTCAGTGCTGTGTCCG; rDNA-F4: ACCGCCACCTCCTTGACCTGAGTC; rDNA-R4: AAACGACTCAGGTCAAGGAGGTGG; rDNA-F5: ACCGTCGCTCTGTCACCCAGGCTG; rDNA-R5: AAACCAGCCTGGGTGACAGAGCGA; rDNA-F6: ACCGTCTTCGGTAGCTGGGATTAC; rDNA-R6: AAACGTAATCCCAGCTACCGAAGA; rDNA-F7: ACCGTCCTGGGCCTCCCAAAGTGC; rDNA-R7: AAACGCACTTTGGGAGGCCCAGGA; rDNA-F8: ACCGTAGGCCATTGCACTGTAGCC; rDNA-R8: AAACGGCTACAGTGCAATGGCCTA; rDNA-F9: ACCGACGTCTGTCATCCCGAGGTC; rDNA-R9: AAACGACCTCGGGATGACAGACGT; rDNA-F10: ACCGCGAAATGGAGTCAGGCGCCG; rDNA-R10: AAACCGGCGCCTGACTCCATTTCG; rDNA-F11: ACCGTGTAACCCCAGCTACTCGGG; rDNA-R11: AAACCCCGAGTAGCTGGGGTTACA; rDNA-F12: ACCGAATTGCTTGAACCTGGCAGG; rDNA-R12: AAACCCTGCCAGGTTCAAGCAATT; rDNA-F13: ACCGCTGGGCGACAGAGTGAGACC; rDNA-R13: AAACGGTCTCACTCTGTCGCCCAG; rDNA-F14: ACCGACTGTATTGCTACTGGGCTA; rDNA-R14: AAACTAGCCCAGTAGCAATACAGT; rDNA-F15: ACCGCCGAGAGGAATCTAGACAGG; rDNA-R15: AAACCCTGTCTAGATTCCTCTCGG; rDNA-F16: ACCGGGTCCCCGCTTGGATGCGAG; rDNA-R16: AAACCTCGCATCCAAGCGGGGACC. HeLa cells were seeded onto 6 well plates. siRNAs were transfected as described above a day after seeding. After another 24 hours, the dCas9 and gsRNAs were transfected to corresponding wells with Lipofectamine 2000 according to manufacturer’s instruction at a concentration of 200 ng of dCas9 and 750 ng sgRNA individually. Signals were evaluated 42 hours post-transfection using Nickon fluorescence microscope. Actinomycin D was applied at a concentration of 0.04 ug/ml for 3 hours at 42 hours post-CRISPR construct transfection.

## Supplemental materials

They include additional supporting data including 1 table and figures.

## Acknowledgements

This study was funded by NIH grants 2 R01 GM078555-09 and U01CA260699 to S.H., 1R35GM131687 to S.J.B and U01 DA-040588 to T.P. and Paul Kaufman, UMass Chan Medical School, the latter part of the NIH 4-D Nucleome Initiative. We thank Lihua Julie Zhu and Paul Kaufman at UMass Chan Medical School for expertise on the bioinformatics search of GC-rich loci in the human genome. We also thank Dr. Samuel Sondalle for part of protein expression analyses. In addition, we thank Ulrike Kutay (Institute of Biochemistry, ETH Zurich, Zurich, Switzerland) for the GFP-RPL29 construct, Zehra Nizami and Joseph Gall (Department of Embryology, Carnegie Institution for Science) for the anti-FLASH antibody. We would like to thank Tom Volpe, Northwestern U. for critical reading of the manuscript.

## Author contributions

S.H. designed the project; H.M. and T.P. provided the CRISPR-dCas9 rDNA labeling system; C.W.; S.J.B. provided the UTP4 siRNA knockdown plasmid and contributed to the manuscript preparation; S.H. and T.P. wrote the paper with input from all authors.

## Supplemental material

**Suppl. Table 1:**
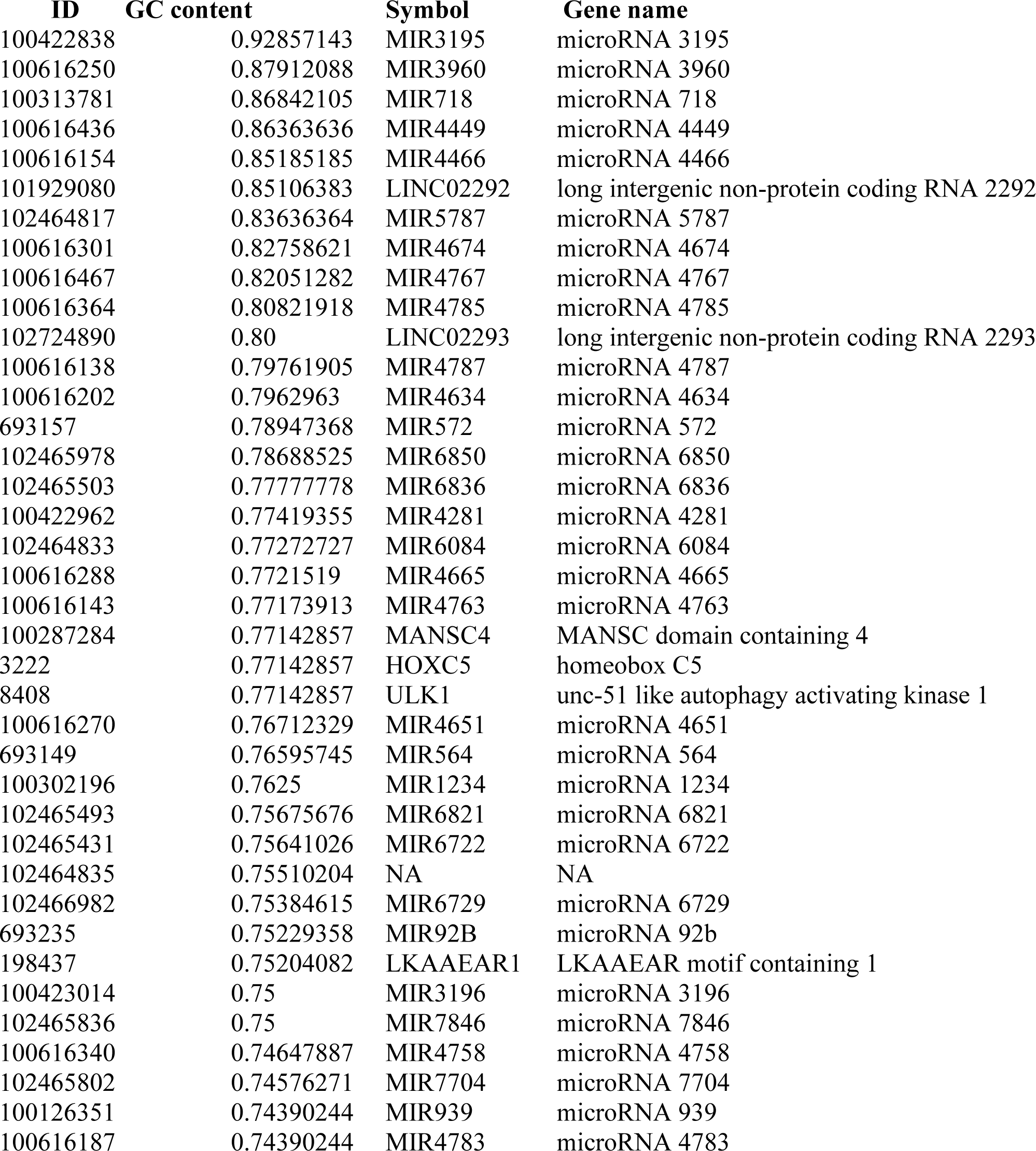
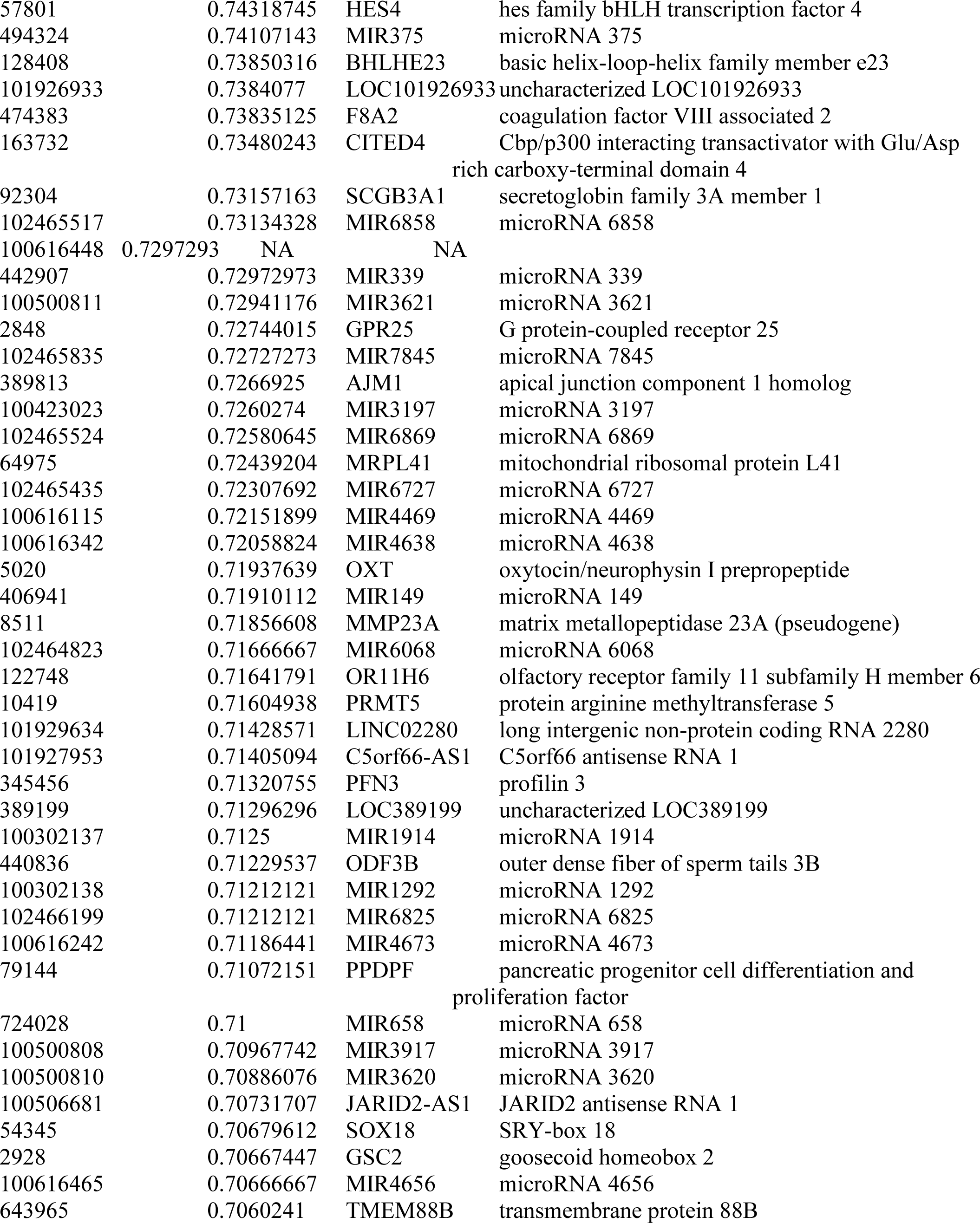
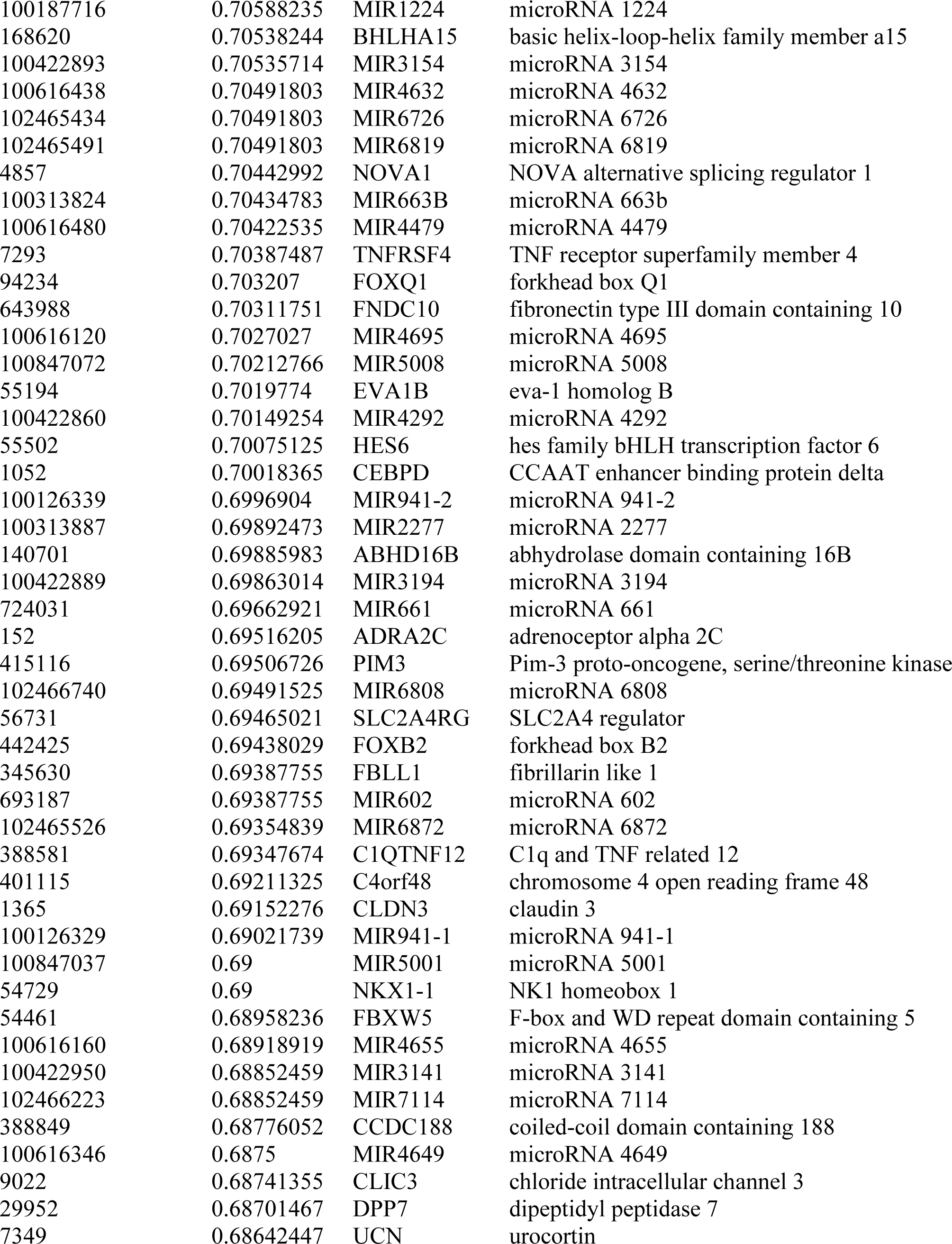
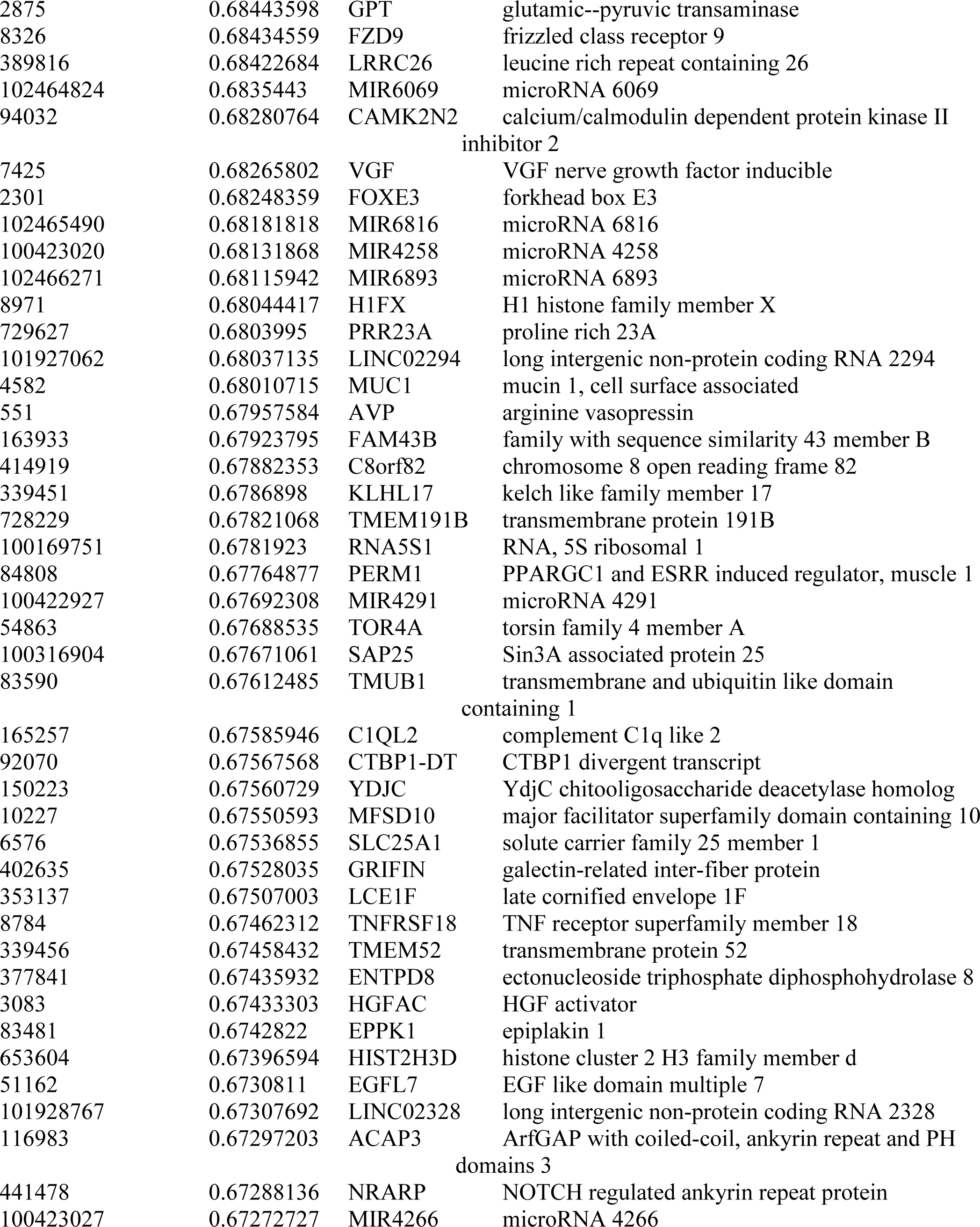
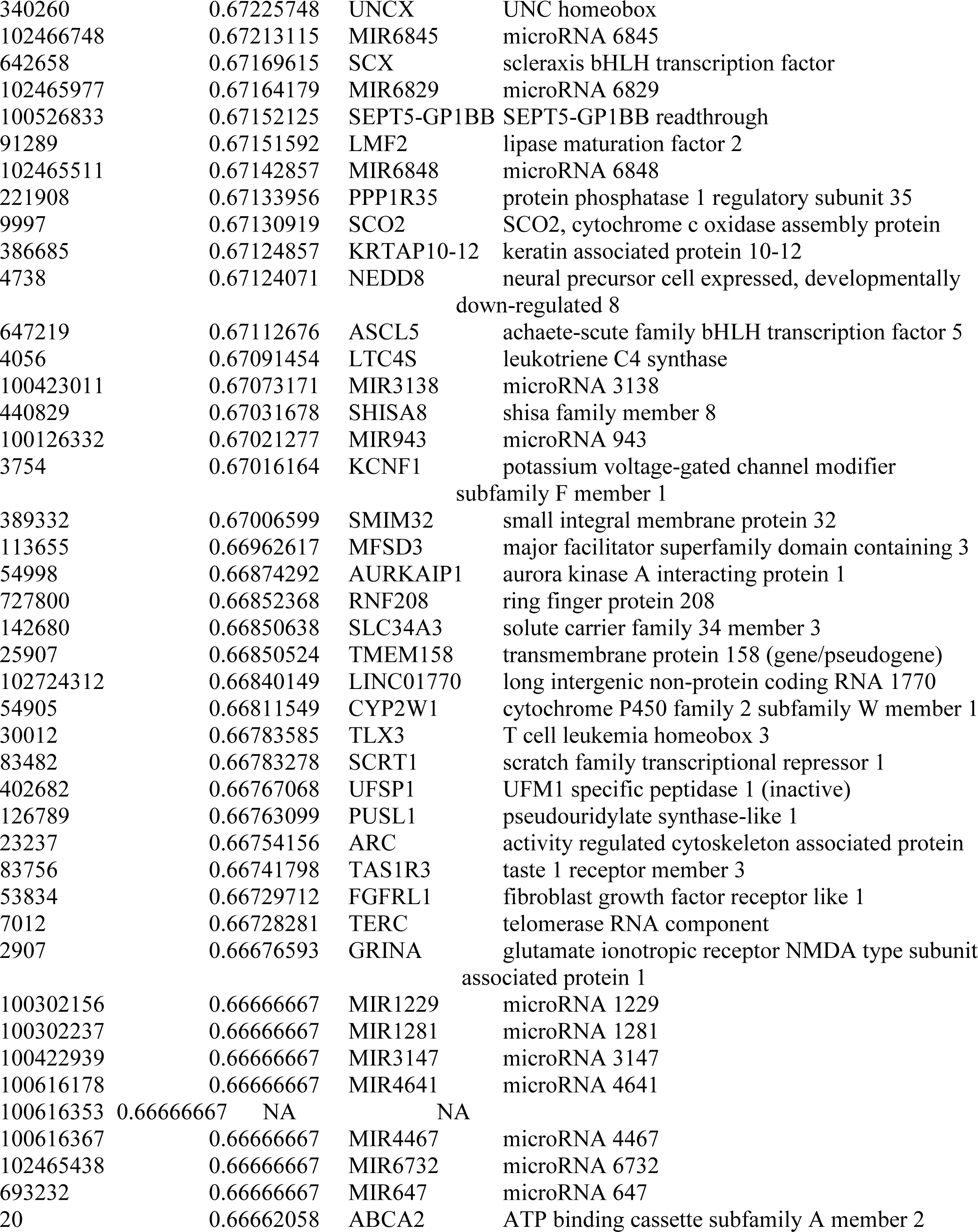
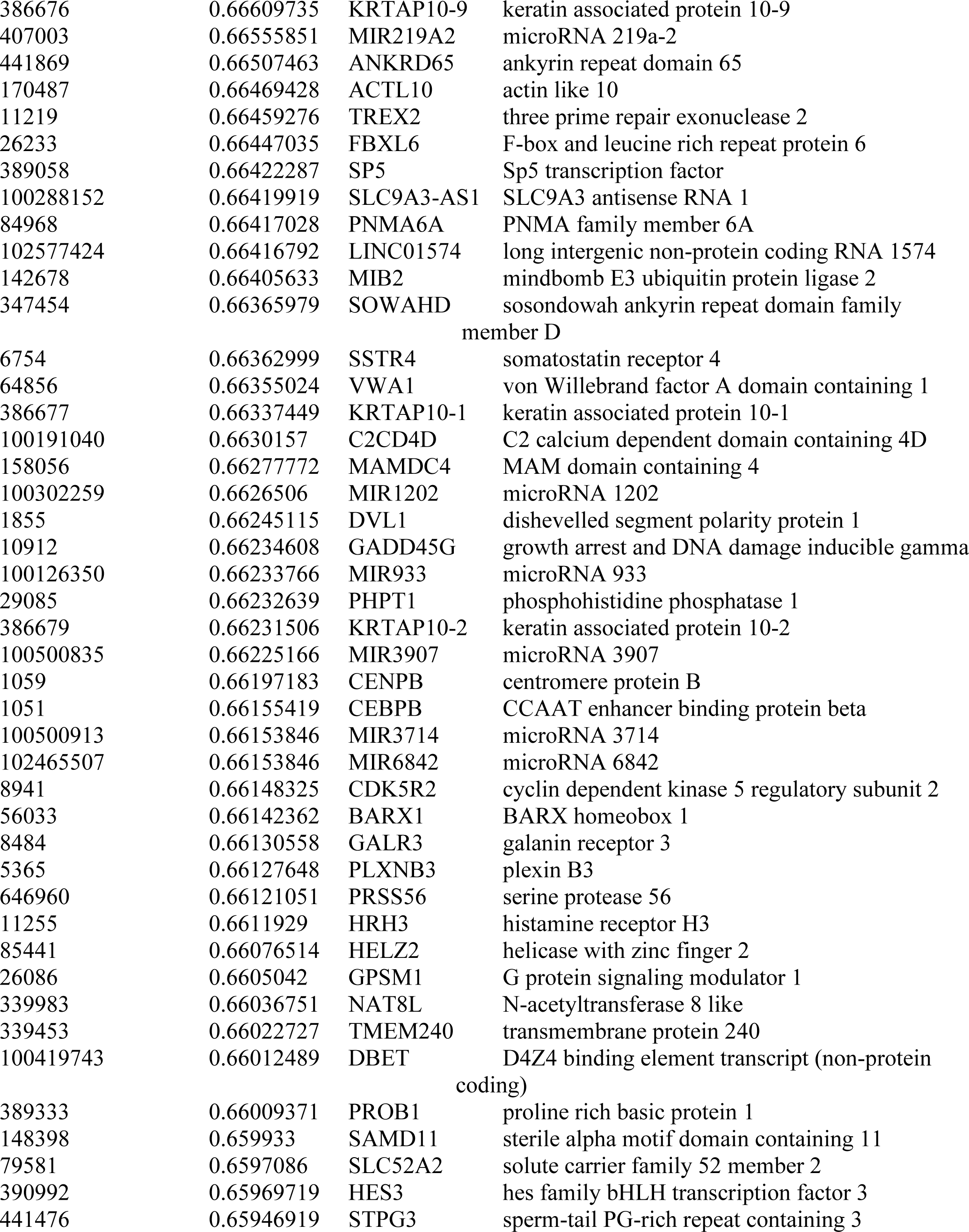
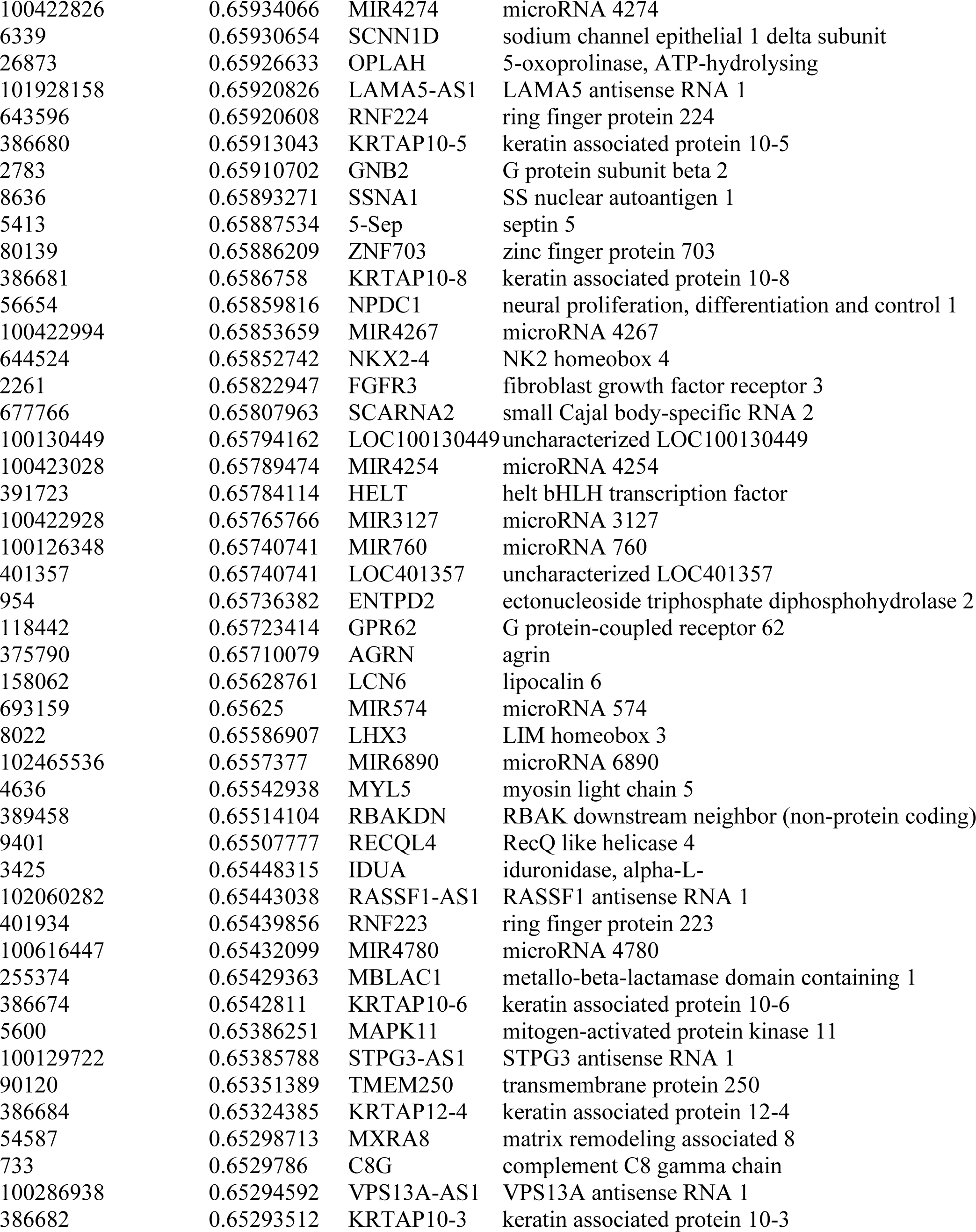
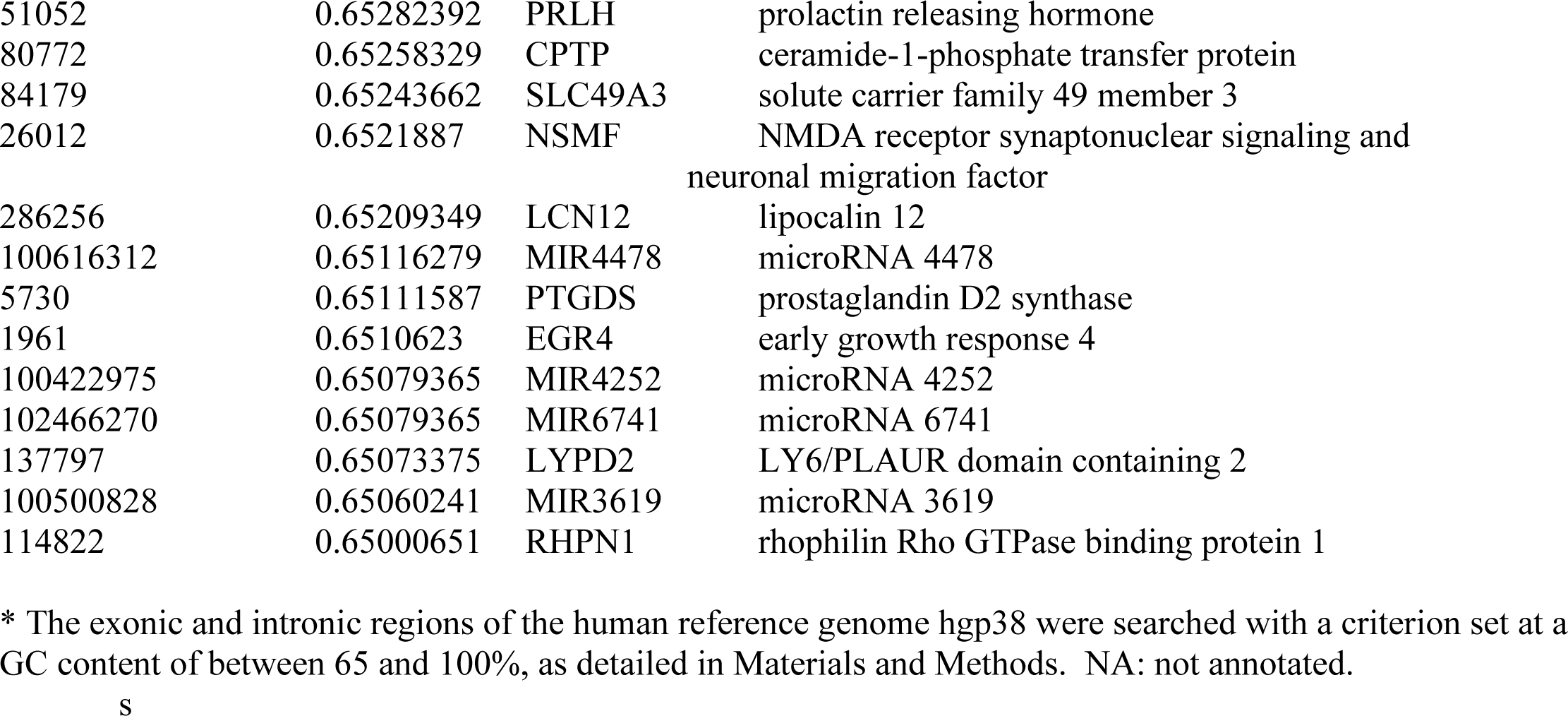
* The exonic and intronic regions of the human reference genome hgp38 were searched with a criterion set at a GC content of between 65 and 100%, as detailed in Materials and Methods. NA: not annotated.

| **ID** | **GC content** | **Symbol** | **Gene name** |
| --- | --- | --- | --- |
| 100422838 | 0.92857143 | MIR3195 | microRNA 3195 |
| 100616250 | 0.87912088 | MIR3960 | microRNA 3960 |
| 100313781 | 0.86842105 | MIR718 | microRNA 718 |
| 100616436 | 0.86363636 | MIR4449 | microRNA 4449 |
| 100616154 | 0.85185185 | MIR4466 | microRNA 4466 |
| 101929080 | 0.85106383 | LINC02292 | long intergenic non-protein coding RNA 2292 |
| 102464817 | 0.83636364 | MIR5787 | microRNA 5787 |
| 100616301 | 0.82758621 | MIR4674 | microRNA 4674 |
| 100616467 | 0.82051282 | MIR4767 | microRNA 4767 |
| 100616364 | 0.80821918 | MIR4785 | microRNA 4785 |
| 102724890 | 0.80 | LINC02293 | long intergenic non-protein coding RNA 2293 |
| 100616138 | 0.79761905 | MIR4787 | microRNA 4787 |
| 100616202 | 0.7962963 | MIR4634 | microRNA 4634 |
| 693157 | 0.78947368 | MIR572 | microRNA 572 |
| 102465978 | 0.78688525 | MIR6850 | microRNA 6850 |
| 102465503 | 0.77777778 | MIR6836 | microRNA 6836 |
| 100422962 | 0.77419355 | MIR4281 | microRNA 4281 |
| 102464833 | 0.77272727 | MIR6084 | microRNA 6084 |
| 100616288 | 0.7721519 | MIR4665 | microRNA 4665 |
| 100616143 | 0.77173913 | MIR4763 | microRNA 4763 |
| 100287284 | 0.77142857 | MANSC4 | MANSC domain containing 4 |
| 3222 | 0.77142857 | HOXC5 | homeobox C5 |
| 8408 | 0.77142857 | ULK1 | unc-51 like autophagy activating kinase 1 |
| 100616270 | 0.76712329 | MIR4651 | microRNA 4651 |
| 693149 | 0.76595745 | MIR564 | microRNA 564 |
| 100302196 | 0.7625 | MIR1234 | microRNA 1234 |
| 102465493 | 0.75675676 | MIR6821 | microRNA 6821 |
| 102465431 | 0.75641026 | MIR6722 | microRNA 6722 |
| 102464835 | 0.75510204 | NA | NA |
| 102466982 | 0.75384615 | MIR6729 | microRNA 6729 |
| 693235 | 0.75229358 | MIR92B | microRNA 92b |
| 198437 | 0.75204082 | LKAAEAR1 | LKAAEAR motif containing 1 |
| 100423014 | 0.75 | MIR3196 | microRNA 3196 |
| 102465836 | 0.75 | MIR7846 | microRNA 7846 |
| 100616340 | 0.74647887 | MIR4758 | microRNA 4758 |
| 102465802 | 0.74576271 | MIR7704 | microRNA 7704 |
| 100126351 | 0.74390244 | MIR939 | microRNA 939 |
| 100616187 | 0.74390244 | MIR4783 | microRNA 4783 |
| 57801 | 0.74318745 | HES4 | hes family bHLH transcription factor 4 |
| 494324 | 0.74107143 | MIR375 | microRNA 375 |
| 128408 | 0.73850316 | BHLHE23 | basic helix-loop-helix family member e23 |
| 101926933 | 0.7384077 | LOC101926933 | uncharacterized LOC101926933 |
| 474383 | 0.73835125 | F8A2 | coagulation factor VIII associated 2 |
| 163732 | 0.73480243 | CITED4 | Cbp/p300 interacting transactivator with Glu/Asp<br>rich carboxy-terminal domain 4 |
| 92304 | 0.73157163 | SCGB3A1 | secretoglobin family 3A member 1 |
| 102465517 | 0.73134328 | MIR6858 | microRNA 6858 |
| 100616448 | 0.7297293 | NA | NA |
| 442907 | 0.72972973 | MIR339 | microRNA 339 |
| 100500811 | 0.72941176 | MIR3621 | microRNA 3621 |
| 2848 | 0.72744015 | GPR25 | G protein-coupled receptor 25 |
| 102465835 | 0.72727273 | MIR7845 | microRNA 7845 |
| 389813 | 0.7266925 | AJM1 | apical junction component 1 homolog |
| 100423023 | 0.7260274 | MIR3197 | microRNA 3197 |
| 102465524 | 0.72580645 | MIR6869 | microRNA 6869 |
| 64975 | 0.72439204 | MRPL41 | mitochondrial ribosomal protein L41 |
| 102465435 | 0.72307692 | MIR6727 | microRNA 6727 |
| 100616115 | 0.72151899 | MIR4469 | microRNA 4469 |
| 100616342 | 0.72058824 | MIR4638 | microRNA 4638 |
| 5020 | 0.71937639 | OXT | oxytocin/neurophysin I prepropeptide |
| 406941 | 0.71910112 | MIR149 | microRNA 149 |
| 8511 | 0.71856608 | MMP23A | matrix metalloproteinase 23A (pseudogene) |
| 102464823 | 0.71666667 | MIR6068 | microRNA 6068 |
| 122748 | 0.71641791 | OR11H6 | olfactory receptor family 11 subfamily H member 6 |
| 10419 | 0.71604938 | PRMT5 | protein arginine methyltransferase 5 |
| 101929634 | 0.71428571 | LINC02280 | long intergenic non-protein coding RNA 2280 |
| 101927953 | 0.71405094 | C5orf66-AS1 | C5orf66 antisense RNA 1 |
| 345456 | 0.71320755 | PFN3 | profilin 3 |
| 389199 | 0.71296296 | LOC389199 | uncharacterized LOC389199 |
| 100302137 | 0.7125 | MIR1914 | microRNA 1914 |
| 440836 | 0.71229537 | ODF3B | outer dense fiber of sperm tails 3B |
| 100302138 | 0.71212121 | MIR1292 | microRNA 1292 |
| 102466199 | 0.71212121 | MIR6825 | microRNA 6825 |
| 100616242 | 0.71186441 | MIR4673 | microRNA 4673 |
| 79144 | 0.71072151 | PPDPF | pancreatic progenitor cell differentiation and<br>proliferation factor |
| 724028 | 0.71 | MIR658 | microRNA 658 |
| 100500808 | 0.70967742 | MIR3917 | microRNA 3917 |
| 100500810 | 0.70886076 | MIR3620 | microRNA 3620 |
| 100506681 | 0.70731707 | JARID2-AS1 | JARID2 antisense RNA 1 |
| 54345 | 0.70679612 | SOX18 | SRY-box 18 |
| 2928 | 0.70667447 | GSC2 | goosecoid homeobox 2 |
| 100616465 | 0.70666667 | MIR4656 | microRNA 4656 |
| 643965 | 0.7060241 | TMEM88B | transmembrane protein 88B |
| 100187716 | 0.70588235 | MIR1224 | microRNA 1224 |
| 168620 | 0.70538244 | BHLHA15 | basic helix-loop-helix family member a15 |
| 100422893 | 0.70535714 | MIR3154 | microRNA 3154 |
| 100616438 | 0.70491803 | MIR4632 | microRNA 4632 |
| 102465434 | 0.70491803 | MIR6726 | microRNA 6726 |
| 102465491 | 0.70491803 | MIR6819 | microRNA 6819 |
| 4857 | 0.70442992 | NOVA1 | NOVA alternative splicing regulator 1 |
| 100313824 | 0.70434783 | MIR663B | microRNA 663b |
| 100616480 | 0.70422535 | MIR4479 | microRNA 4479 |
| 7293 | 0.70387487 | TNFRSF4 | TNF receptor superfamily member 4 |
| 94234 | 0.703207 | FOXQ1 | forkhead box Q1 |
| 643988 | 0.70311751 | FNDC10 | fibronectin type III domain containing 10 |
| 100616120 | 0.7027027 | MIR4695 | microRNA 4695 |
| 100847072 | 0.70212766 | MIR5008 | microRNA 5008 |
| 55194 | 0.7019774 | EVA1B | eva-1 homolog B |
| 100422860 | 0.70149254 | MIR4292 | microRNA 4292 |
| 55502 | 0.70075125 | HES6 | hes family bHLH transcription factor 6 |
| 1052 | 0.70018365 | CEBPD | CCAAT enhancer binding protein delta |
| 100126339 | 0.6996904 | MIR941-2 | microRNA 941-2 |
| 100313887 | 0.69892473 | MIR2277 | microRNA 2277 |
| 140701 | 0.69885983 | ABHD16B | abhydrolase domain containing 16B |
| 100422889 | 0.69863014 | MIR3194 | microRNA 3194 |
| 724031 | 0.69662921 | MIR661 | microRNA 661 |
| 152 | 0.69516205 | ADRA2C | adrenoceptor alpha 2C |
| 415116 | 0.69506726 | PIM3 | Pim-3 proto-oncogene, serine/threonine kinase |
| 102466740 | 0.69491525 | MIR6808 | microRNA 6808 |
| 56731 | 0.69465021 | SLC2A4RG | SLC2A4 regulator |
| 442425 | 0.69438029 | FOXB2 | forkhead box B2 |
| 345630 | 0.69387755 | FBLL1 | fibrillarlin like 1 |
| 693187 | 0.69387755 | MIR602 | microRNA 602 |
| 102465526 | 0.69354839 | MIR6872 | microRNA 6872 |
| 388581 | 0.69347674 | C1QTNF12 | C1q and TNF related 12 |
| 401115 | 0.69211325 | C4orf48 | chromosome 4 open reading frame 48 |
| 1365 | 0.69152276 | CLDN3 | claudin 3 |
| 100126329 | 0.69021739 | MIR941-1 | microRNA 941-1 |
| 100847037 | 0.69 | MIR5001 | microRNA 5001 |
| 54729 | 0.69 | NKX1-1 | NK1 homeobox 1 |
| 54461 | 0.68958236 | FBXW5 | F-box and WD repeat domain containing 5 |
| 100616160 | 0.68918919 | MIR4655 | microRNA 4655 |
| 100422950 | 0.68852459 | MIR3141 | microRNA 3141 |
| 102466223 | 0.68852459 | MIR7114 | microRNA 7114 |
| 388849 | 0.68776052 | CCDC188 | coiled-coil domain containing 188 |
| 100616346 | 0.6875 | MIR4649 | microRNA 4649 |
| 9022 | 0.68741355 | CLIC3 | chloride intracellular channel 3 |
| 29952 | 0.68701467 | DPP7 | dipeptidyl peptidase 7 |
| 7349 | 0.68642447 | UCN | urocortin |
| 2875 | 0.68443598 | GPT | glutamic--pyruvic transaminase |
| 8326 | 0.68434559 | FZD9 | frizzled class receptor 9 |
| 389816 | 0.68422684 | LRRC26 | leucine rich repeat containing 26 |
| 102464824 | 0.6835443 | MIR6069 | microRNA 6069 |
| 94032 | 0.68280764 | CAMK2N2 | calcium/calmodulin dependent protein kinase II inhibitor 2 |
| 7425 | 0.68265802 | VGF | VGF nerve growth factor inducible |
| 2301 | 0.68248359 | FOXE3 | forkhead box E3 |
| 102465490 | 0.68181818 | MIR6816 | microRNA 6816 |
| 100423020 | 0.68131868 | MIR4258 | microRNA 4258 |
| 102466271 | 0.68115942 | MIR6893 | microRNA 6893 |
| 8971 | 0.68044417 | H1FX | H1 histone family member X |
| 729627 | 0.6803995 | PRR23A | proline rich 23A |
| 101927062 | 0.68037135 | LINC02294 | long intergenic non-protein coding RNA 2294 |
| 4582 | 0.68010715 | MUC1 | mucin 1, cell surface associated |
| 551 | 0.67957584 | AVP | arginine vasopressin |
| 163933 | 0.67923795 | FAM43B | family with sequence similarity 43 member B |
| 414919 | 0.67882353 | C8orf82 | chromosome 8 open reading frame 82 |
| 339451 | 0.6786898 | KLHL17 | kelch like family member 17 |
| 728229 | 0.67821068 | TMEM191B | transmembrane protein 191B |
| 100169751 | 0.6781923 | RNA5S1 | RNA, 5S ribosomal 1 |
| 84808 | 0.67764877 | PERM1 | PPARGC1 and ESRR induced regulator, muscle 1 |
| 100422927 | 0.67692308 | MIR4291 | microRNA 4291 |
| 54863 | 0.67688535 | TOR4A | torsin family 4 member A |
| 100316904 | 0.67671061 | SAP25 | Sin3A associated protein 25 |
| 83590 | 0.67612485 | TMUB1 | transmembrane and ubiquitin like domain containing 1 |
| 165257 | 0.67585946 | C1QL2 | complement C1q like 2 |
| 92070 | 0.67567568 | CTBP1-DT | CTBP1 divergent transcript |
| 150223 | 0.67560729 | YDJC | YdjC chitoooligosaccharide deacetylase homolog |
| 10227 | 0.67550593 | MFSD10 | major facilitator superfamily domain containing 10 |
| 6576 | 0.67536855 | SLC25A1 | solute carrier family 25 member 1 |
| 402635 | 0.67528035 | GRIFIN | galectin-related inter-fiber protein |
| 353137 | 0.67507003 | LCE1F | late cornified envelope 1F |
| 8784 | 0.67462312 | TNFRSF18 | TNF receptor superfamily member 18 |
| 339456 | 0.67458432 | TMEM52 | transmembrane protein 52 |
| 377841 | 0.67435932 | ENTPD8 | ectonucleoside triphosphate diphosphohydrolase 8 |
| 3083 | 0.67433303 | HGFAC | HGF activator |
| 83481 | 0.6742822 | EPPK1 | epiplakin 1 |
| 653604 | 0.67396594 | HIST2H3D | histone cluster 2 H3 family member d |
| 51162 | 0.6730811 | EGFL7 | EGF like domain multiple 7 |
| 101928767 | 0.67307692 | LINC02328 | long intergenic non-protein coding RNA 2328 |
| 116983 | 0.67297203 | ACAP3 | ArfGAP with coiled-coil, ankyrin repeat and PH domains 3 |
| 441478 | 0.67288136 | NRARP | NOTCH regulated ankyrin repeat protein |
| 100423027 | 0.67272727 | MIR4266 | microRNA 4266 |
| 340260 | 0.67225748 | UNCX | UNC homeobox |
| 102466748 | 0.67213115 | MIR6845 | microRNA 6845 |
| 642658 | 0.67169615 | SCX | scleraxis bHLH transcription factor |
| 102465977 | 0.67164179 | MIR6829 | microRNA 6829 |
| 100526833 | 0.67152125 | SEPT5-GP1BB | SEPT5-GP1BB readthrough |
| 91289 | 0.67151592 | LMF2 | lipase maturation factor 2 |
| 102465511 | 0.67142857 | MIR6848 | microRNA 6848 |
| 221908 | 0.67133956 | PPP1R35 | protein phosphatase 1 regulatory subunit 35 |
| 9997 | 0.67130919 | SCO2 | SCO2, cytochrome c oxidase assembly protein |
| 386685 | 0.67124857 | KRTAP10-12 | keratin associated protein 10-12 |
| 4738 | 0.67124071 | NEDD8 | neural precursor cell expressed, developmentally down-regulated 8 |
| 647219 | 0.67112676 | ASCL5 | achaete-scute family bHLH transcription factor 5 |
| 4056 | 0.67091454 | LTC4S | leukotriene C4 synthase |
| 100423011 | 0.67073171 | MIR3138 | microRNA 3138 |
| 440829 | 0.67031678 | SHISA8 | shisa family member 8 |
| 100126332 | 0.67021277 | MIR943 | microRNA 943 |
| 3754 | 0.67016164 | KCNF1 | potassium voltage-gated channel modifier subfamily F member 1 |
| 389332 | 0.67006599 | SMIM32 | small integral membrane protein 32 |
| 113655 | 0.66962617 | MFSD3 | major facilitator superfamily domain containing 3 |
| 54998 | 0.66874292 | AURKAIP1 | aurora kinase A interacting protein 1 |
| 727800 | 0.66852368 | RNF208 | ring finger protein 208 |
| 142680 | 0.66850638 | SLC34A3 | solute carrier family 34 member 3 |
| 25907 | 0.66850524 | TMEM158 | transmembrane protein 158 (gene/pseudogene) |
| 102724312 | 0.66840149 | LINC01770 | long intergenic non-protein coding RNA 1770 |
| 54905 | 0.66811549 | CYP2W1 | cytochrome P450 family 2 subfamily W member 1 |
| 30012 | 0.66783585 | TLX3 | T cell leukemia homeobox 3 |
| 83482 | 0.66783278 | SCRT1 | scratch family transcriptional repressor 1 |
| 402682 | 0.66767068 | UFSP1 | UFM1 specific peptidase 1 (inactive) |
| 126789 | 0.66763099 | PUSL1 | pseudouridylate synthase-like 1 |
| 23237 | 0.66754156 | ARC | activity regulated cytoskeleton associated protein |
| 83756 | 0.66741798 | TAS1R3 | taste 1 receptor member 3 |
| 53834 | 0.66729712 | FGFRL1 | fibroblast growth factor receptor like 1 |
| 7012 | 0.66728281 | TERC | telomerase RNA component |
| 2907 | 0.66676593 | GRINA | glutamate ionotropic receptor NMDA type subunit associated protein 1 |
| 100302156 | 0.66666667 | MIR1229 | microRNA 1229 |
| 100302237 | 0.66666667 | MIR1281 | microRNA 1281 |
| 100422939 | 0.66666667 | MIR3147 | microRNA 3147 |
| 100616178 | 0.66666667 | MIR4641 | microRNA 4641 |
| 100616353 | 0.66666667 | NA | NA |
| 100616367 | 0.66666667 | MIR4467 | microRNA 4467 |
| 102465438 | 0.66666667 | MIR6732 | microRNA 6732 |
| 693232 | 0.66666667 | MIR647 | microRNA 647 |
| 20 | 0.66662058 | ABCA2 | ATP binding cassette subfamily A member 2 |
| 386676 | 0.66609735 | KRTAP10-9 | keratin associated protein 10-9 |
| 407003 | 0.66555851 | MIR219A2 | microRNA 219a-2 |
| 441869 | 0.66507463 | ANKRD65 | ankyrin repeat domain 65 |
| 170487 | 0.66469428 | ACTL10 | actin like 10 |
| 11219 | 0.66459276 | TREX2 | three prime repair exonuclease 2 |
| 26233 | 0.66447035 | FBXL6 | F-box and leucine rich repeat protein 6 |
| 389058 | 0.66422287 | SP5 | Sp5 transcription factor |
| 100288152 | 0.66419919 | SLC9A3-AS1 | SLC9A3 antisense RNA 1 |
| 84968 | 0.66417028 | PNMA6A | PNMA family member 6A |
| 102577424 | 0.66416792 | LINC01574 | long intergenic non-protein coding RNA 1574 |
| 142678 | 0.66405633 | MIB2 | mindbomb E3 ubiquitin protein ligase 2 |
| 347454 | 0.66365979 | SOWAHD | sosondowah ankyrin repeat domain family member D |
| 6754 | 0.66362999 | SSTR4 | somatostatin receptor 4 |
| 64856 | 0.66355024 | VWA1 | von Willebrand factor A domain containing 1 |
| 386677 | 0.66337449 | KRTAP10-1 | keratin associated protein 10-1 |
| 100191040 | 0.6630157 | C2CD4D | C2 calcium dependent domain containing 4D |
| 158056 | 0.66277772 | MAMDC4 | MAM domain containing 4 |
| 100302259 | 0.6626506 | MIR1202 | microRNA 1202 |
| 1855 | 0.66245115 | DVL1 | dishevelled segment polarity protein 1 |
| 10912 | 0.66234608 | GADD45G | growth arrest and DNA damage inducible gamma |
| 100126350 | 0.66233766 | MIR933 | microRNA 933 |
| 29085 | 0.66232639 | PHPT1 | phosphohistidine phosphatase 1 |
| 386679 | 0.66231506 | KRTAP10-2 | keratin associated protein 10-2 |
| 100500835 | 0.66225166 | MIR3907 | microRNA 3907 |
| 1059 | 0.66197183 | CENPB | centromere protein B |
| 1051 | 0.66155419 | CEBPB | CCAAT enhancer binding protein beta |
| 100500913 | 0.66153846 | MIR3714 | microRNA 3714 |
| 102465507 | 0.66153846 | MIR6842 | microRNA 6842 |
| 8941 | 0.66148325 | CDK5R2 | cyclin dependent kinase 5 regulatory subunit 2 |
| 56033 | 0.66142362 | BARX1 | BARX homeobox 1 |
| 8484 | 0.66130558 | GALR3 | galanin receptor 3 |
| 5365 | 0.66127648 | PLXNB3 | plexin B3 |
| 646960 | 0.66121051 | PRSS56 | serine protease 56 |
| 11255 | 0.6611929 | HRH3 | histamine receptor H3 |
| 85441 | 0.66076514 | HELZ2 | helicase with zinc finger 2 |
| 26086 | 0.6605042 | GPSM1 | G protein signaling modulator 1 |
| 339983 | 0.66036751 | NAT8L | N-acetyltransferase 8 like |
| 339453 | 0.66022727 | TMEM240 | transmembrane protein 240 |
| 100419743 | 0.66012489 | DBET | D4Z4 binding element transcript (non-protein coding) |
| 389333 | 0.66009371 | PROB1 | proline rich basic protein 1 |
| 148398 | 0.659933 | SAMD11 | sterile alpha motif domain containing 11 |
| 79581 | 0.6597086 | SLC52A2 | solute carrier family 52 member 2 |
| 390992 | 0.65969719 | HES3 | hes family bHLH transcription factor 3 |
| 441476 | 0.65946919 | STPG3 | sperm-tail PG-rich repeat containing 3 |
| 100422826 | 0.65934066 | MIR4274 | microRNA 4274 |
| 6339 | 0.65930654 | SCNN1D | sodium channel epithelial 1 delta subunit |
| 26873 | 0.65926633 | OPLAH | 5-oxoprolinase, ATP-hydrolysing |
| 101928158 | 0.65920826 | LAMA5-AS1 | LAMA5 antisense RNA 1 |
| 643596 | 0.65920608 | RNF224 | ring finger protein 224 |
| 386680 | 0.65913043 | KRTAP10-5 | keratin associated protein 10-5 |
| 2783 | 0.65910702 | GNB2 | G protein subunit beta 2 |
| 8636 | 0.65893271 | SSNA1 | SS nuclear autoantigen 1 |
| 5413 | 0.65887534 | 5-Sep | septin 5 |
| 80139 | 0.65886209 | ZNF703 | zinc finger protein 703 |
| 386681 | 0.6586758 | KRTAP10-8 | keratin associated protein 10-8 |
| 56654 | 0.65859816 | NPDC1 | neural proliferation, differentiation and control 1 |
| 100422994 | 0.65853659 | MIR4267 | microRNA 4267 |
| 644524 | 0.65852742 | NKX2-4 | NK2 homeobox 4 |
| 2261 | 0.65822947 | FGFR3 | fibroblast growth factor receptor 3 |
| 677766 | 0.65807963 | SCARNA2 | small Cajal body-specific RNA 2 |
| 100130449 | 0.65794162 | LOC100130449 | uncharacterized LOC100130449 |
| 100423028 | 0.65789474 | MIR4254 | microRNA 4254 |
| 391723 | 0.65784114 | HELT | helt bHLH transcription factor |
| 100422928 | 0.65765766 | MIR3127 | microRNA 3127 |
| 100126348 | 0.65740741 | MIR760 | microRNA 760 |
| 401357 | 0.65740741 | LOC401357 | uncharacterized LOC401357 |
| 954 | 0.65736382 | ENTPD2 | ectonucleoside triphosphate diphosphohydrolase 2 |
| 118442 | 0.65723414 | GPR62 | G protein-coupled receptor 62 |
| 375790 | 0.65710079 | AGRN | agrin |
| 158062 | 0.65628761 | LCN6 | lipocalin 6 |
| 693159 | 0.65625 | MIR574 | microRNA 574 |
| 8022 | 0.65586907 | LHX3 | LIM homeobox 3 |
| 102465536 | 0.6557377 | MIR6890 | microRNA 6890 |
| 4636 | 0.65542938 | MYL5 | myosin light chain 5 |
| 389458 | 0.65514104 | RBAKDN | RBAK downstream neighbor (non-protein coding) |
| 9401 | 0.65507777 | RECQL4 | RecQ like helicase 4 |
| 3425 | 0.65448315 | IDUA | iduronidase, alpha-L- |
| 102060282 | 0.65443038 | RASSF1-AS1 | RASSF1 antisense RNA 1 |
| 401934 | 0.65439856 | RNF223 | ring finger protein 223 |
| 100616447 | 0.65432099 | MIR4780 | microRNA 4780 |
| 255374 | 0.65429363 | MBLAC1 | metallo-beta-lactamase domain containing 1 |
| 386674 | 0.6542811 | KRTAP10-6 | keratin associated protein 10-6 |
| 5600 | 0.65386251 | MAPK11 | mitogen-activated protein kinase 11 |
| 100129722 | 0.65385788 | STPG3-AS1 | STPG3 antisense RNA 1 |
| 90120 | 0.65351389 | TMEM250 | transmembrane protein 250 |
| 386684 | 0.65324385 | KRTAP12-4 | keratin associated protein 12-4 |
| 54587 | 0.65298713 | MXRA8 | matrix remodeling associated 8 |
| 733 | 0.6529786 | C8G | complement C8 gamma chain |
| 100286938 | 0.65294592 | VPS13A-AS1 | VPS13A antisense RNA 1 |
| 386682 | 0.65293512 | KRTAP10-3 | keratin associated protein 10-3 |
| 51052 | 0.65282392 | PRLH | prolactin releasing hormone |
| 80772 | 0.65258329 | CPTP | ceramide-1-phosphate transfer protein |
| 84179 | 0.65243662 | SLC49A3 | solute carrier family 49 member 3 |
| 26012 | 0.6521887 | NSMF | NMDA receptor synaptonuclear signaling and neuronal migration factor |
| 286256 | 0.65209349 | LCN12 | lipocalin 12 |
| 100616312 | 0.65116279 | MIR4478 | microRNA 4478 |
| 5730 | 0.65111587 | PTGDS | prostaglandin D2 synthase |
| 1961 | 0.6510623 | EGR4 | early growth response 4 |
| 100422975 | 0.65079365 | MIR4252 | microRNA 4252 |
| 102466270 | 0.65079365 | MIR6741 | microRNA 6741 |
| 137797 | 0.65073375 | LYPD2 | LY6/PLAUR domain containing 2 |
| 100500828 | 0.65060241 | MIR3619 | microRNA 3619 |
| 114822 | 0.65000651 | RHPN1 | rhophilin Rho GTPase binding protein 1 |

**Suppl. Figure 1:**
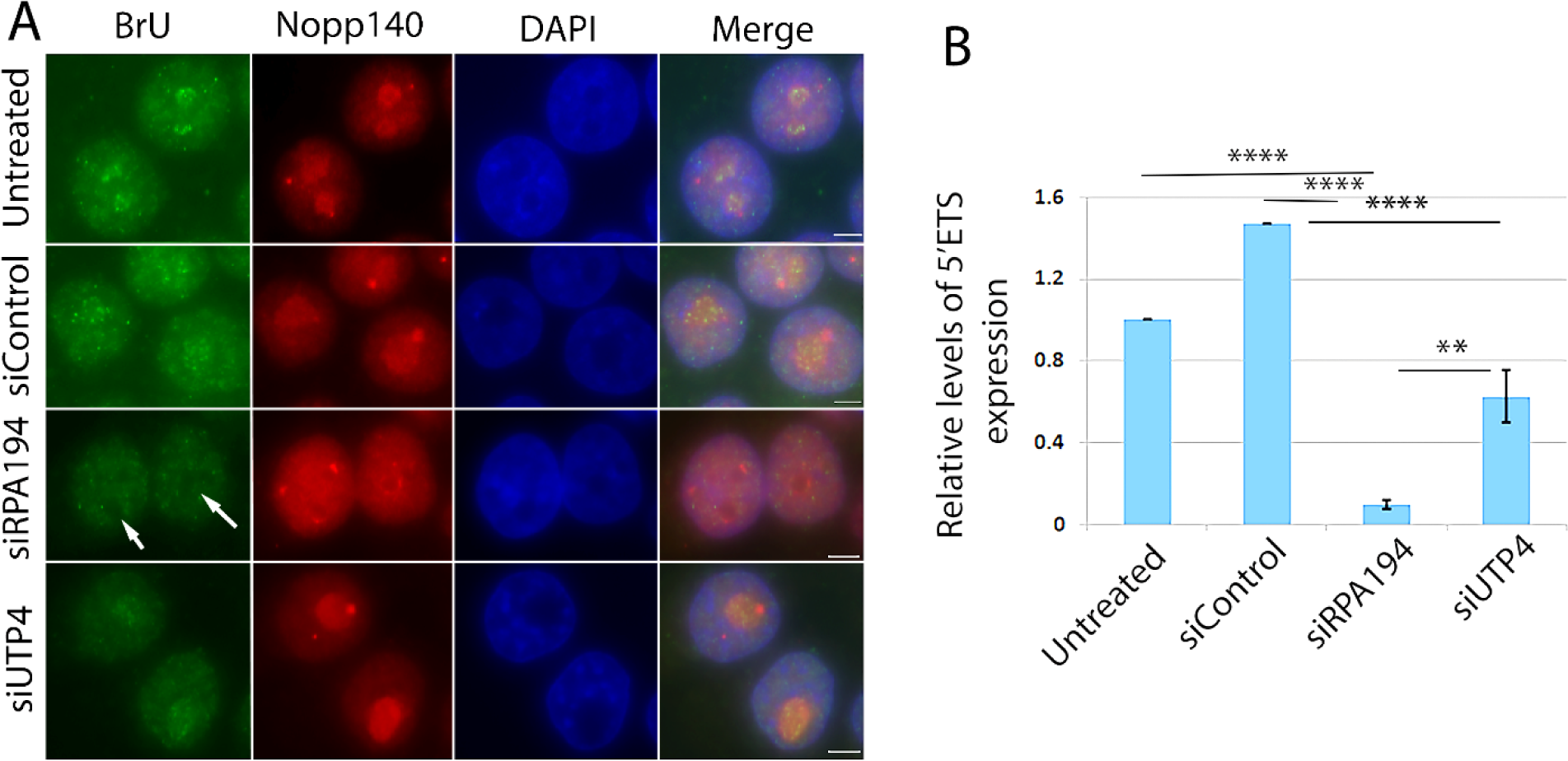
The transcription of rDNA was significantly reduced in cells treated with siRPA194 and to a lesser extent with siUTP4. (A) BrU pulse labeling for 5 minutes showed a significant loss of BrU signal in siRPA194 treated cells (A, arrows), compared to controls where more intense labeling was detected in nucleoli. (B) Quantitative RT-PCR showed a significant reduction in 5’ETS RNA expression in siRPA194 and siUTP4 treated cells. Bar= 5 mm. ****p<0.001, **p<0.01

**Suppl. Figure 2:**
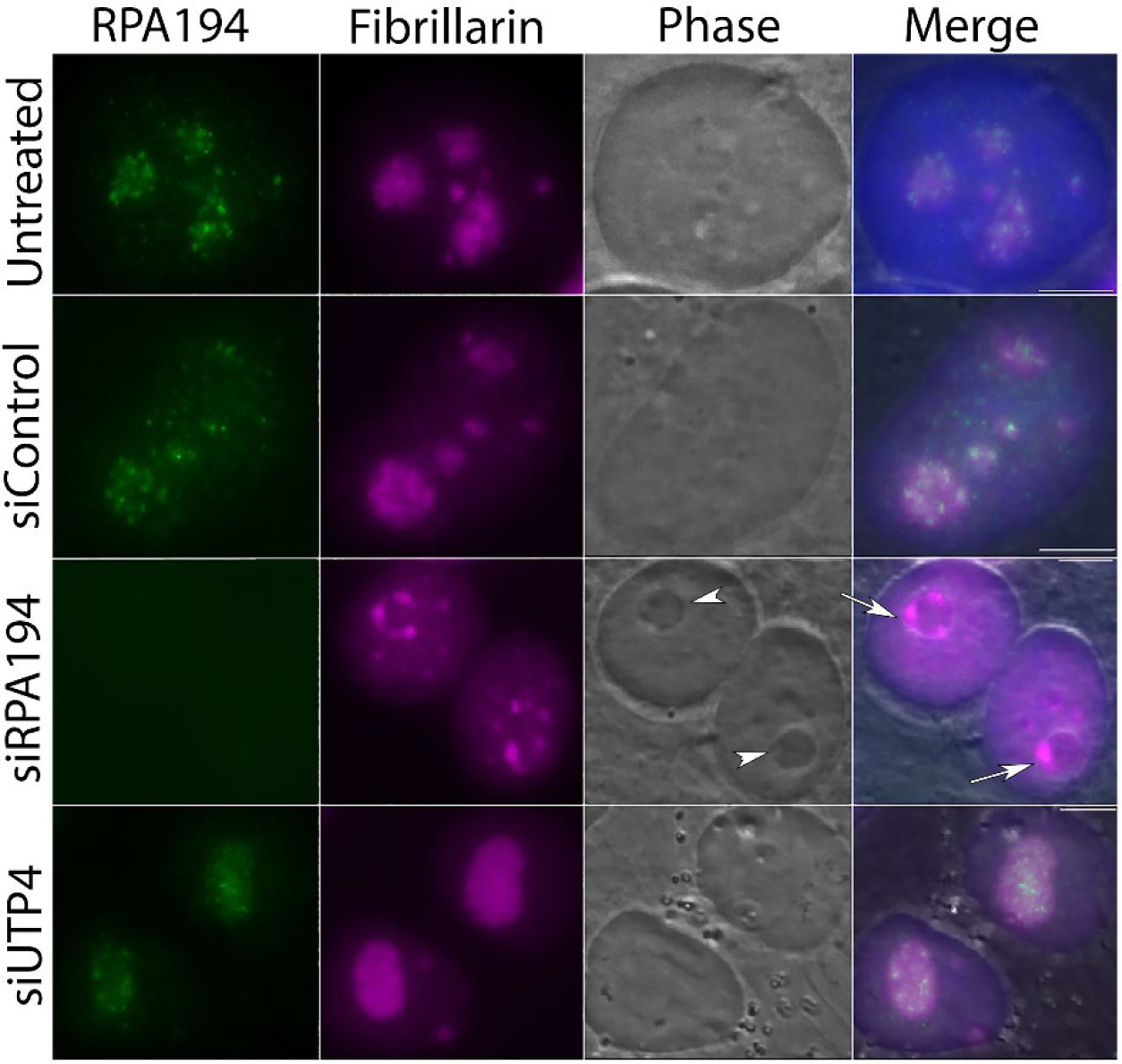
siRPA194, but not siUTP4 treated HeLa cells, lost the RPA194 signal (green). Nucleolar segregation was detected by the immunolabeling of fibrillarin (magenta). Cap-like structures enriched with firbrillar components (arrows) juxtaposed the reminants of the nucleoli primarily containing granular components (arrowheads). Bars= 5 mm

**Suppl. Figure 3:**
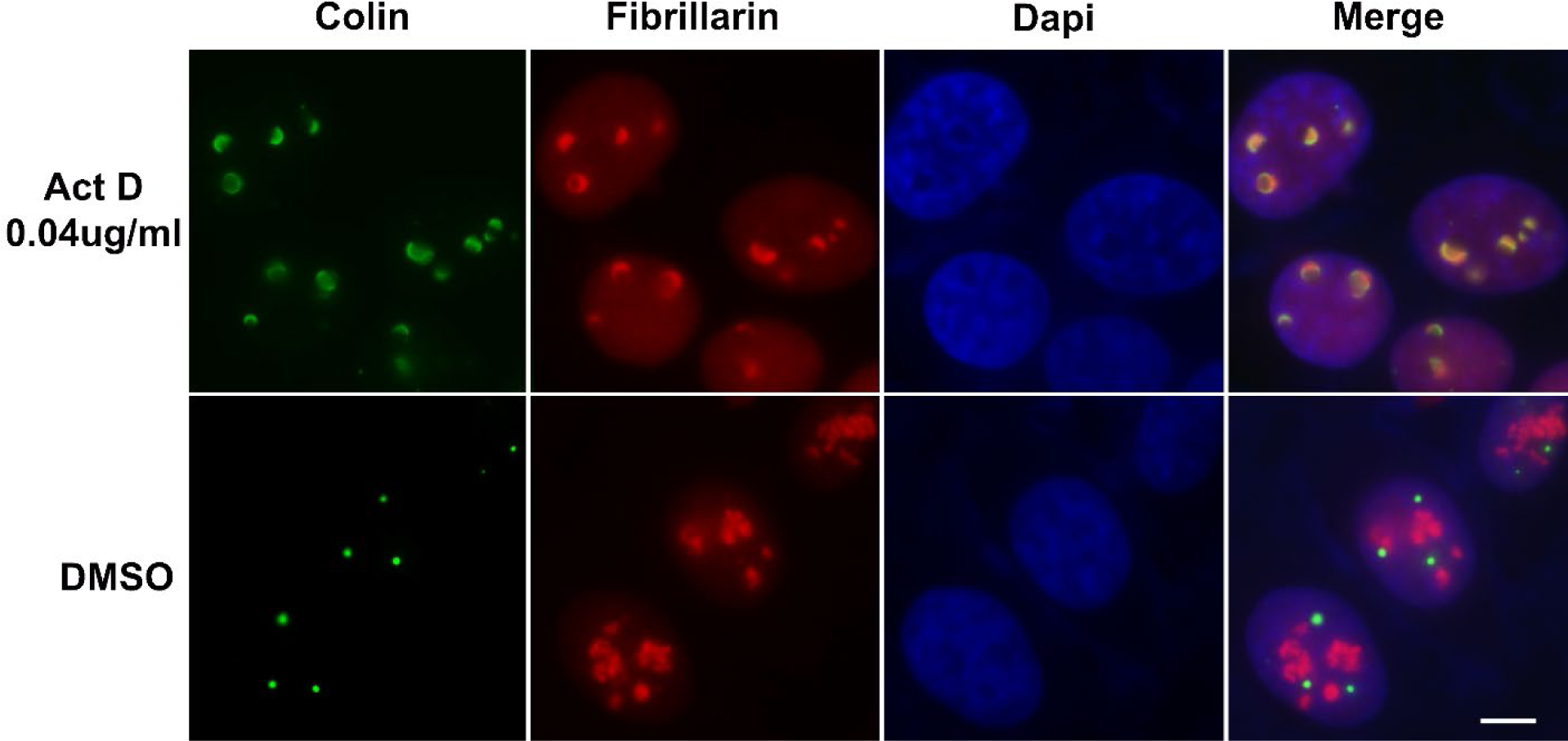
Treatment with a low concentration (0.04ug/ml) of actinomycin D (Act D) induces nucleolar segregations and disruption of Cajal bodies as detected by anti-fibrillarin antibodies (red) and anti-coilin (green) antibodies. Bar= 5 mm

**Suppl. Figure 4:**
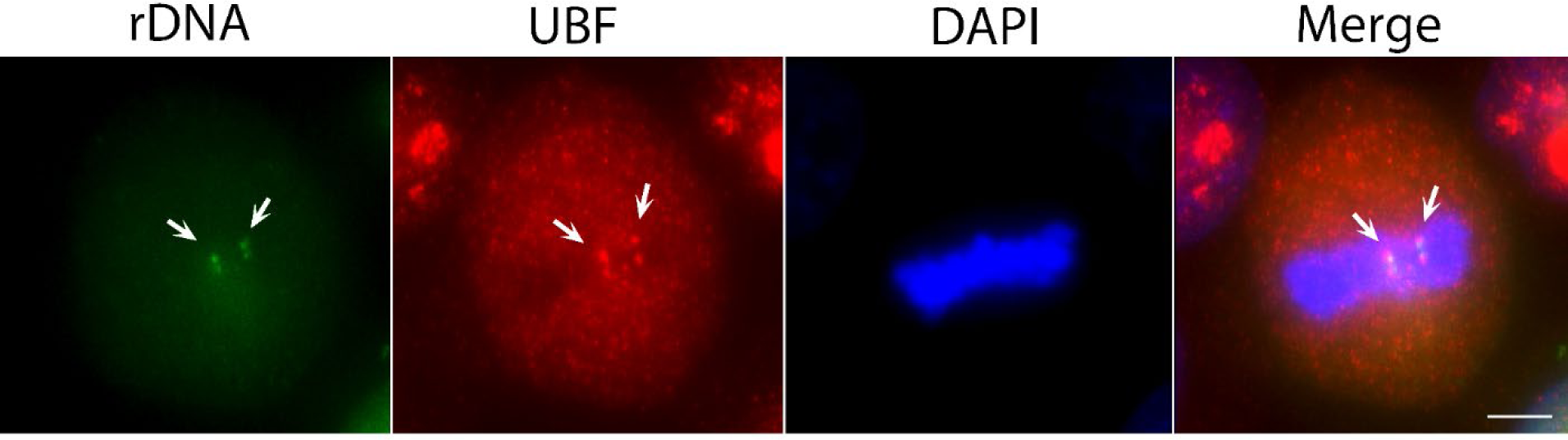
The CRISPR-Cas9 sgRNA labeling of rDNA was validated using mitotic cells where UBF was found to be colocalized with the labeling (arrows), confirming that the sgRNA sets indeed labeled rDNA chromatin on mitotic chromosomes (arrow). Bar= 5 mm

**Suppl. Fig. 5:**
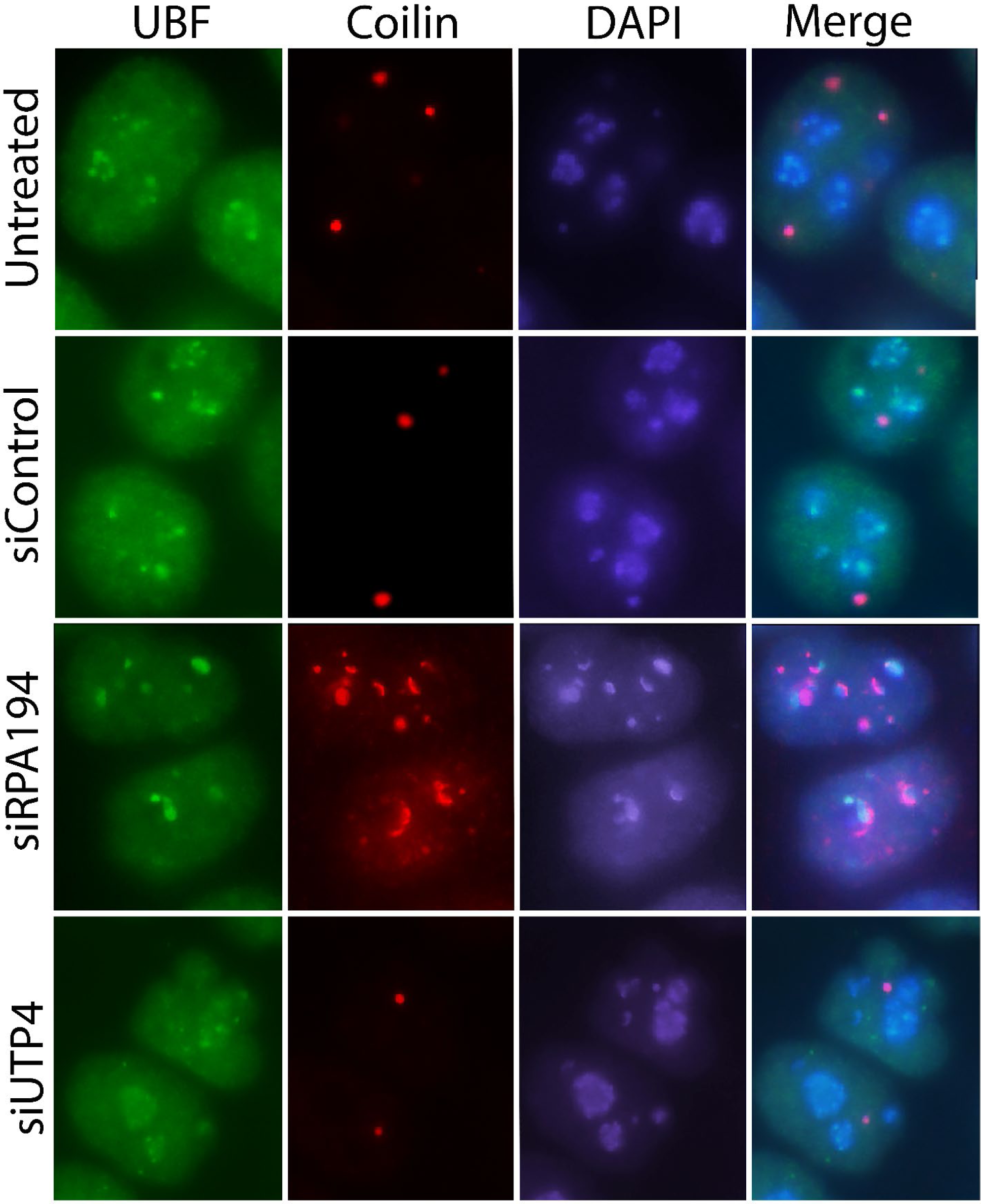
Cajal bodies are significantly altered in siRPA194 treated cells. Cells were co-immunolabeled with antibodies against UBF and coilin. Bar= 5 mm

**Suppl. Figure 6:**
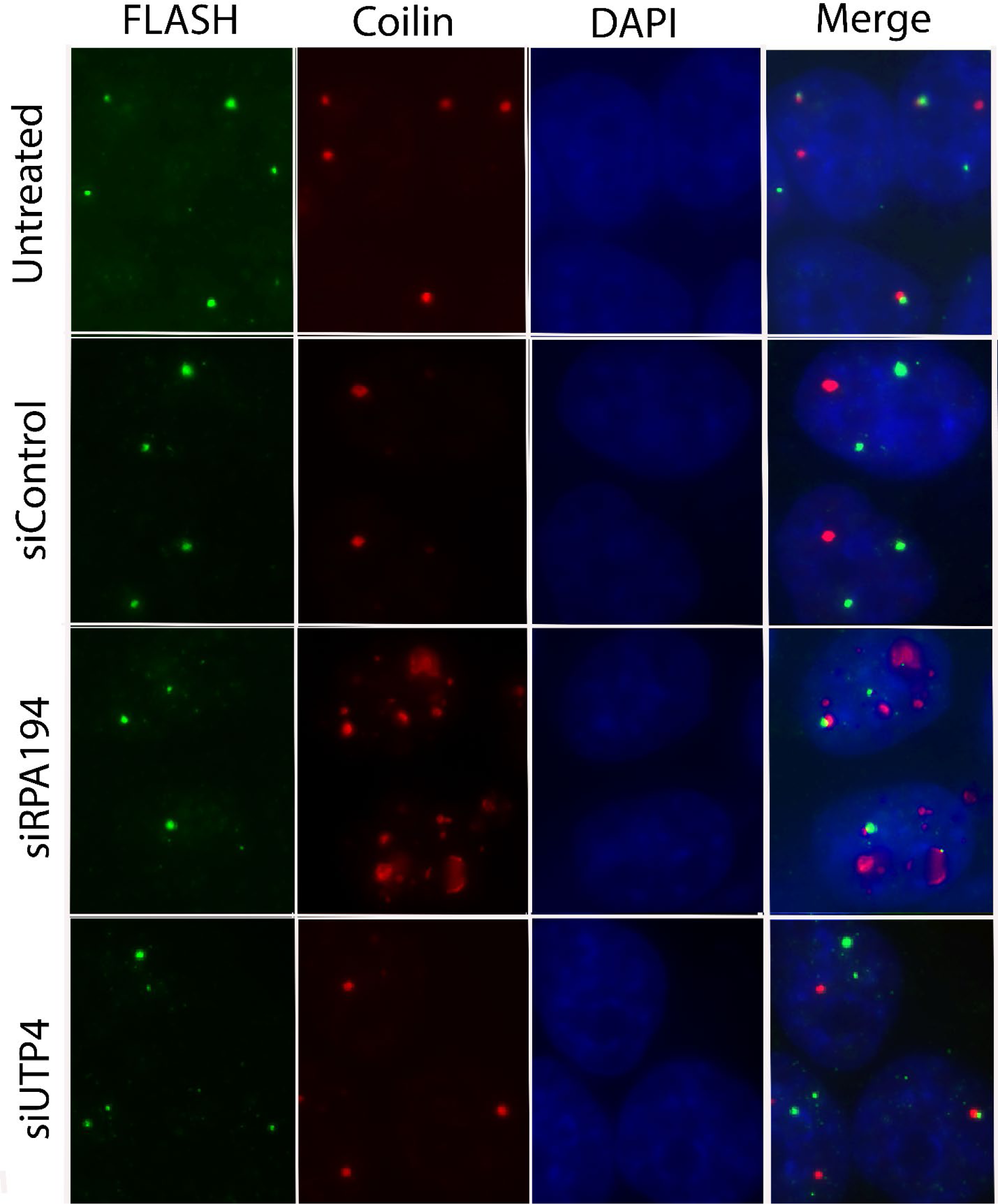
While Cajal bodies change significantly in siRPA194 treated cells, histone locus bodies do not show detectable changes. Bar= 5 mm

## Abbreviations

PML: promyelocytic leukaemia
BrU: Bromouridine
EM: Electron microscope
CRISPR: Clustered regularly interspaced short palindromic repeats
NOR: Nucleolar organization region
PNC: Perinucleolar compartment
FC: Fibrillar center
DFC: Dense fibrillar components
GC: Granular components

## Notes

### Competing Interest Statement

The authors have declared no competing interest.

